# Predicting Pathway Dependency in Head and Neck Squamous Cell Carcinoma from Hallmark Transcriptomic Features

**DOI:** 10.64898/2026.10.26.761775

**Authors:** Yashmin Afshar, Ali Goli

**Author notes:** Corresponding author: Dr. Yashmin Afshar, DDS. Address: Tehran University of Medical Sciences, Children’s Medical Center Hospital, Dr. Qarib St, Keshavarz Blvd, Tehran 14194, Iran.

## Abstract

Despite frequent pathway activation, targeted molecular therapy remains limited in head and neck squamous cell carcinoma (HNSCC), highlighting the need to distinguish activation from dependence. Dependency is functional, context-specific, and coupled; mutations or expression alone cannot establish it. No validated framework identifies tumor pathway dependencies from baseline molecular profiles.

We developed a transcriptome-based framework to infer pathway dependency in HNSCC. Nine Elastic Net models used 50 MSigDB Hallmark activity profiles from 953 GDSC cell lines and drug responses pooled across compounds annotated to each pathway. Cross-validated R² ranged from 0.17 (IGF1R) to 0.48 (P53); leave-one-pathway-out validation retained rank agreement (Spearman ρ = 0.19–0.56) for pathways with no drug response observed during training. We validated the models using pharmacological, genomic, and proteomic data, then applied them to TCGA-HNSC (n = 565) and GSE65858 (n = 256), examining HPV status, age, stage, tobacco history, and CD8+ infiltration.

Seven of nine pathways were higher in HPV+ tumors and replicated directionally in GSE65858. Among TP53-wild-type tumors, HPV remained linked to greater P53 dependency (β = +0.37, q = 0.010); HPV+ tumors also had higher p53 transcriptional activity, suggesting residual p53 function despite E6 expression. The codependency signature had a dominant global axis (PC1 = 62.3% variance; Cronbach’s α = 0.91), replicated across cohorts (Tucker congruence = 0.996), but largely reflected CD8⁺ T-cell infiltration (ρ = 0.80).

The framework uses RNA-sequenced bulk tumor cohorts to assess baseline transcriptome-based dependencies and match pathway-directed agents to likely responders before treatment. It is not prognostic.

**Author Summary:** Cancers may have active pathways with abundant protein expression but not rely on them for survival. Distinguishing pathway activation from pathway dependence, therefore, can improve molecular targeted therapy outcomes and guide combination therapies. Capturing dependency patterns in head and neck squamous cell carcinomas becomes more pressing given the poor outcomes of targeted therapies in cases with well-known mutated proteins, such as PI3K. In this study, we investigated pathway-dependency patterns in head and neck squamous cell carcinomas by training models on pharmacological data and validating them at the genomic and proteomic levels in head and neck squamous cell carcinoma patients.

## 1. Introduction

Head and neck squamous cell carcinoma (HNSCC) (1) is renowned for its genomic heterogeneity, with recurrent mutations in genes that each introduce new biological characteristics to the cancer cell population; for instance, PIK3CA mutations induce radio/chemoresistance in HNSCC, while NOTCH1 mutations create a dependency on the PI3K pathway (2–4). Despite the rising incidence and an overall unfavorable prognosis, cytotoxic chemotherapy and radiotherapy remain the primary therapeutic interventions, with only recent advances from immunotherapy drugs, including pembrolizumab and nivolumab for recurrent/metastatic HNSCC (5–7). Unlike immunotherapy, molecularly targeted approaches have had limited efficacy. Combination therapy with Buparlisib, an oral pan-PI3K inhibitor targeting all 4 isoforms of class I PI3K, and paclitaxel in PD-1/PD-L1-pretreated recurrent/metastatic HNSCC did not improve overall survival compared with paclitaxel alone (8). Moreover, clinical trials of PI3K, AKT, and mTOR inhibitors as monotherapy in HNSCC have shown limited efficacy of 5.3%, 2.8%, and 31%, respectively (9). This is despite the PI3K/AKT/mTOR (PAM) pathway being activated in approximately 90% of HNSCC lesions through diverse mechanisms, including EGFR activation (47%), PIK3CA amplifications and mutations (14.2% and 8.6%), PI3K overexpression (27.2%), and PTEN loss (10-15%) (9, 10). These limited responses further highlight the importance of distinguishing between pathway activation and cancer dependence on a particular pathway.

Dependency is a functional property, not a molecular one, and barely is dependency singular. Pathway dependencies are context-specific and coupled: vulnerability to inhibition of one pathway often depends on the activity of another (11), and inhibiting a dominant pathway frequently triggers compensatory activation of parallel cascades (12).

For next-generation therapies to achieve maximum efficacy and minimize side effects, treatment plans should be more personalized and flexible in response to disease modification. Cancer and the body are closely correlated, and any change in the ecosystem can drive significant shifts over time. This is complicated because the system comprises countless interacting components, and any change can affect the others. Introducing intrinsic or extrinsic stresses, such as starvation, drugs, or hypoxia, can lead to new genetic alterations under stress and subsequently to cancer evolution, making therapies ineffective (13). Moreover, the primary determinant is not genetic alterations; the majority of pathway activation is driven by non-genomic mechanisms, such as upstream receptor signaling, rather than by direct mutation of the pathway’s core components (9, 10). In several cancers, including HNSCC, background diseases also have undeniable effects on the disease’s outcome or even its genetic state. For instance, HNSCC with HPV+ has minimal p53 mutations and a better prognosis, which should be taken into account when prescribing for them (14). The HPV+ background would further introduce a new immune landscape in the body (15). The real challenge is capturing the whole state of the body and disease as an interactive system and defining the treatment plan accordingly. For instance, neglecting inflammation’s role in cancer progression and metastasis may affect treatment outcomes (16, 17). Considering the whole picture, combining cancer-specific therapeutics with drug repurposing may be effective (18). What is missing is not a target but a method for choosing one. No validated framework exists to infer, from a tumor’s baseline molecular profile, which pathways it actually depends on. The full system, however, cannot be captured in a single study, but a baseline dependency readout is a prerequisite for any treatment plan that aims to be adaptive rather than fixed.

Working within the constraints of cell-line-trained, bulk-transcriptomic, and cross-sectional data, we developed a transcriptome-based framework to infer pathway dependency in HNSCC. We used drug-response data from the Genomics of Drug Sensitivity in Cancer (GDSC) project, aggregated LN-IC50 values across all compounds annotated to each pathway to define a functional dependency score, and trained Elastic Net models to predict these scores from the 50 MSigDB Hallmark pathway activity profiles of 953 cell lines. We then validated the results at the pharmacological, genomic, and proteomic levels, applied to TCGA-HNSC to infer dependency in patient tumors, and independently tested in GSE65858. Stratifying by HPV status, we characterized the dependency program in HPV-positive and HPV-negative disease, mapped the co-dependency architecture across the nine pathways, and assessed the prognostic value of the inferred dependencies. Our primary objective is to establish a foundation for selecting an appropriate treatment protocol at the outset of treatment, rather than focusing on prognostication, and to identify which pathways a tumor depends on before therapy, so that pathway-directed agents can be matched to tumors most likely to respond.

## 2. Result

### 2.1 Model validation

To determine whether missing IC50 values were random or associated with pathway dependency, we analyzed missingness across 621 drugs tested on at least 20 cell lines, using the per-drug z-scored scale. Of 188 drug–pathway combinations tested, 31 (16.5%) showed significant missingness after FDR correction; however, missing values per pathway across the nine pathways targeted by our models were extremely small, supporting pathway-wise median imputation (Fig. S1A, Fig.S1B, Table S1).

We trained nine independent ElasticNet regression models to predict pathway-specific drug dependency from 50 Hallmark pathway activity scores in 953 GDSC cell lines, using the negated mean per-drug z-scored LN-IC50 as the target. Cross-validated R² ranged from 0.17 (IGF1R) to 0.48 (P53), with a median of 0.27 across pathways (Table 1, Fig. 1A). Adding pairwise interaction terms among the fifty features yielded ΔR² > 0.01 for IGF1R (ΔR² = +0.020). Based on this screen, we refit the IGF1R pathway with pairwise interactions among its fifty features and retained the remaining pathways in additive form (Fig. 1D, Table S2).

**Figure 1.**
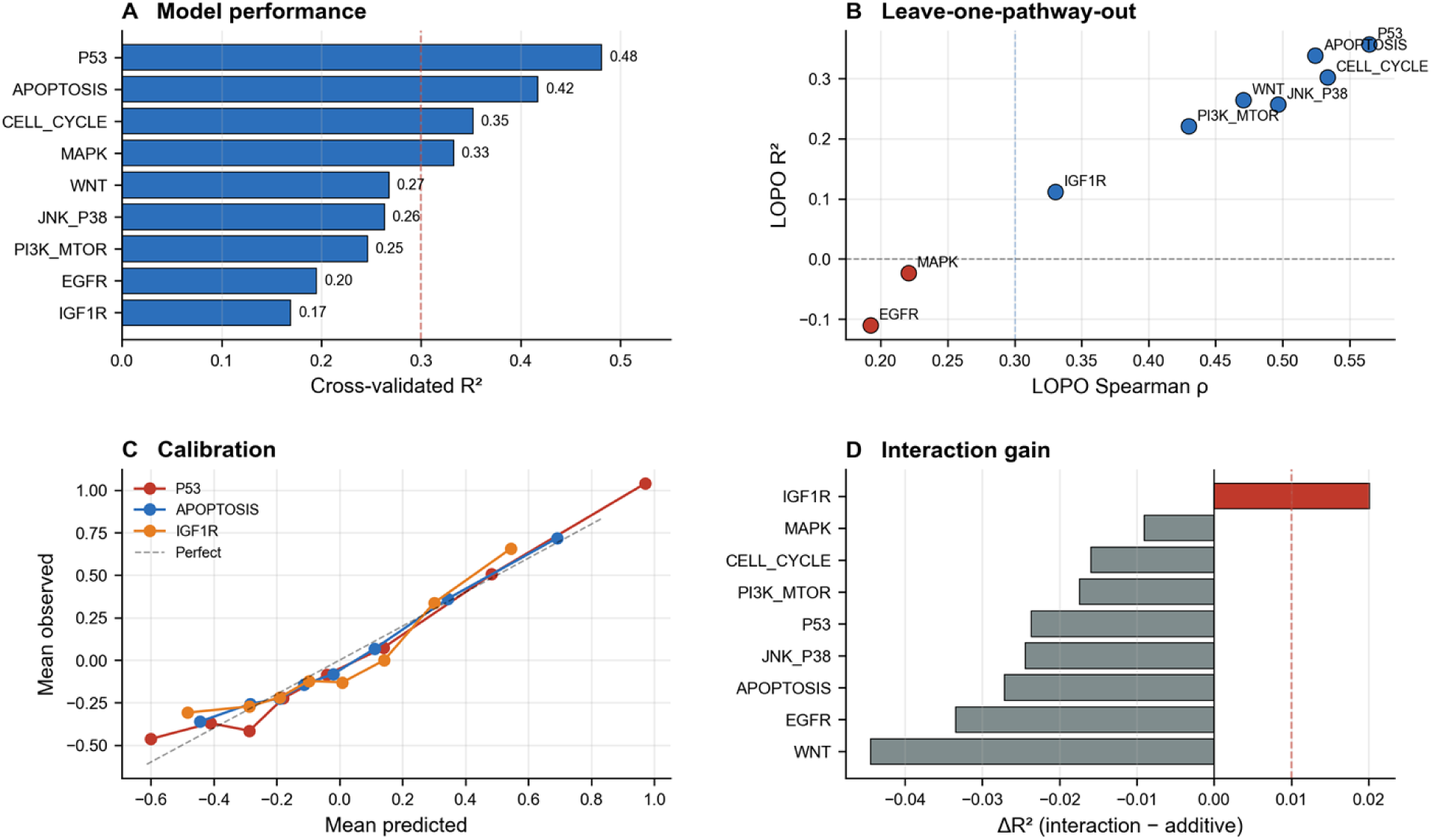
Model training, internal validation, calibration, and interaction screening across nine pathway dependency models. Elastic Net models were trained on 50 Hallmark ssGSEA features to predict drug-derived pathway dependency in 953 GDSC cell lines. (A) Cross-validated R² (5-fold outer CV) for each of the nine pathway models. The dashed red line marks R² = 0.30. (B) Leave-one-pathway-out (LOPO) validation. Each point represents a pathway; the x-axis is the mean Spearman ρ between predicted and held-out dependency scores, and the y-axis is the corresponding LOPO R². Blue points indicate R² ≥ 0; red points indicate R² < 0. (C) Calibration curves for three representative pathways (P53, APOPTOSIS, IGF1R). Out-of-fold predictions were binned into deciles; points show mean predicted vs mean observed dependency per bin, with error bars representing the standard error of the mean. The dashed diagonal is perfect calibration; solid lines are linear fits. (D) Interaction screening. For each pathway, ΔR² = mean R²(interaction model) − mean R²(additive model) under the same 5-fold outer CV. Bars above the red dashed line (ΔR² > 0.01) were retained as interaction models. n = 953 cell lines for all panels. Abbreviations: CV, cross-validation; LOPO, leave-one-pathway-out.

**Table 1.** ElasticNet model performance across nine pathway dependency targets. Base and interaction R² from 5-fold CV on 953 GDSC cell lines; ΔR² > 0.01 retained interactions. LOPO R² and Spearman ρ from leave-one-pathway-out validation. Intercepts are bootstrap 95% CIs (1,000 resamples; 500 for IGF1R).

| Pathway | N drugs | Base $R^2$ | $R^2$ with interactions | $\Delta R^2$ | LOPO $R^2$ | LOPO $\rho$ | Intercept (95% CI) |
| --- | --- | --- | --- | --- | --- | --- | --- |
| P53 | 6 | 0.481 | 0.455 | -0.024 | 0.356 | 0.565 | 0.016 (−0.01–0.05) |
| APOPTOSIS | 23 | 0.417 | 0.388 | -0.027 | 0.338 | 0.524 | 0.019 (−0.00–0.042) |
| CELL_CYCLE | 53 | 0.352 | 0.335 | -0.016 | 0.301 | 0.534 | 0.004 (−0.03–0.04) |
| MAPK | 26 | 0.332 | 0.320 | -0.009 | −0.024 | 0.221 | 0.001 (−0.03–0.04) |
| WNT | 16 | 0.268 | 0.223 | -0.044 | 0.264 | 0.471 | 0.010 (-0.02-0.04) |
| JNK_P38 | 8 | 0.263 | 0.236 | -0.024 | 0.256 | 0.497 | 0.019 (-0.01-0.05) |
| PI3K_MTOR | 59 | 0.246 | 0.235 | -0.017 | 0.220 | 0.430 | 0.009 (-0.02-0.04) |
| EGFR | 12 | 0.195 | 0.163 | -0.033 | -0.111 | 0.193 | 0.010 (-0.03-0.06) |
| IGF1R | 9 | 0.169 | 0.186 | +0.020 | 0.111 | 0.331 | -0.006 (-0.05-0.04) |

To test whether the models captured pathway-level biology rather than memorizing drug–cell line associations, we performed drug-wise hold-out (leave-one-pathway-out) validation. Across all pathways, rank-order agreement between predicted and held-out dependency scores was preserved (LOPO Spearman ρ = 0.19 to 0.56; Table 1), with the strongest agreement for P53 (ρ = 0.56), CELL_CYCLE (ρ = 0.53), and APOPTOSIS (ρ = 0.52) (Fig. 1B, Table 1, Table S3).

To verify that dependencies reflect true drug-sensitivity links, we correlated each drug’s average LN-IC50 with each pathway’s dependency score across 621 drugs and 9 pathways, computing the drug’s own target-pathway dependency LOPO to avoid self-correlation. We identified 1,321 significant negative correlations after FDR correction (ρ < −0.3, q < 0.05, Table S4). The strongest associations were pathway-specific: EGFR dependency was most predictive of EGFR-targeting drugs (Afatinib ρ = −0.82, Sapitinib ρ = −0.76); MAPK dependency was most predictive of MEK/ERK inhibitors (SCH772984 ρ = −0.79, Refametinib ρ = −0.75, Selumetinib ρ = −0.77, ERK_6604 ρ = −0.75); CELL_CYCLE dependency was most predictive of microtubule inhibitors (Vinblastine ρ = −0.80, Vinorelbine ρ = −0.79); IGF1R dependency was most predictive of IGF1R inhibitors (BMS-754807 ρ = −0.73, BMS-536924 ρ = −0.71); PI3K/MTOR dependency was most predictive of PI3K/AKT inhibitors (MK-2206 ρ = −0.72, Pictilisib ρ = −0.68); P53 dependency was most predictive of the Ribociclib (ρ = −0.68); WNT dependency was most predictive of WNT inhibitors (CHIR-99021 ρ = −0.70, SB216763 ρ = −0.63); and APOPTOSIS dependency was most predictive of BCL2 inhibitors (Venetoclax ρ = −0.61, Vinorelbine ρ = −0.65) (Fig. 2A). Binarizing each pathway at the 75th percentile and comparing IC50 between high- and low-dependency cell lines, with the target pathway’s dependency and threshold computed using leave-one-drug-out, yielded 212 significant target validations (q < 0.05; Table S5) (Fig. 2B).

**Figure 2.**
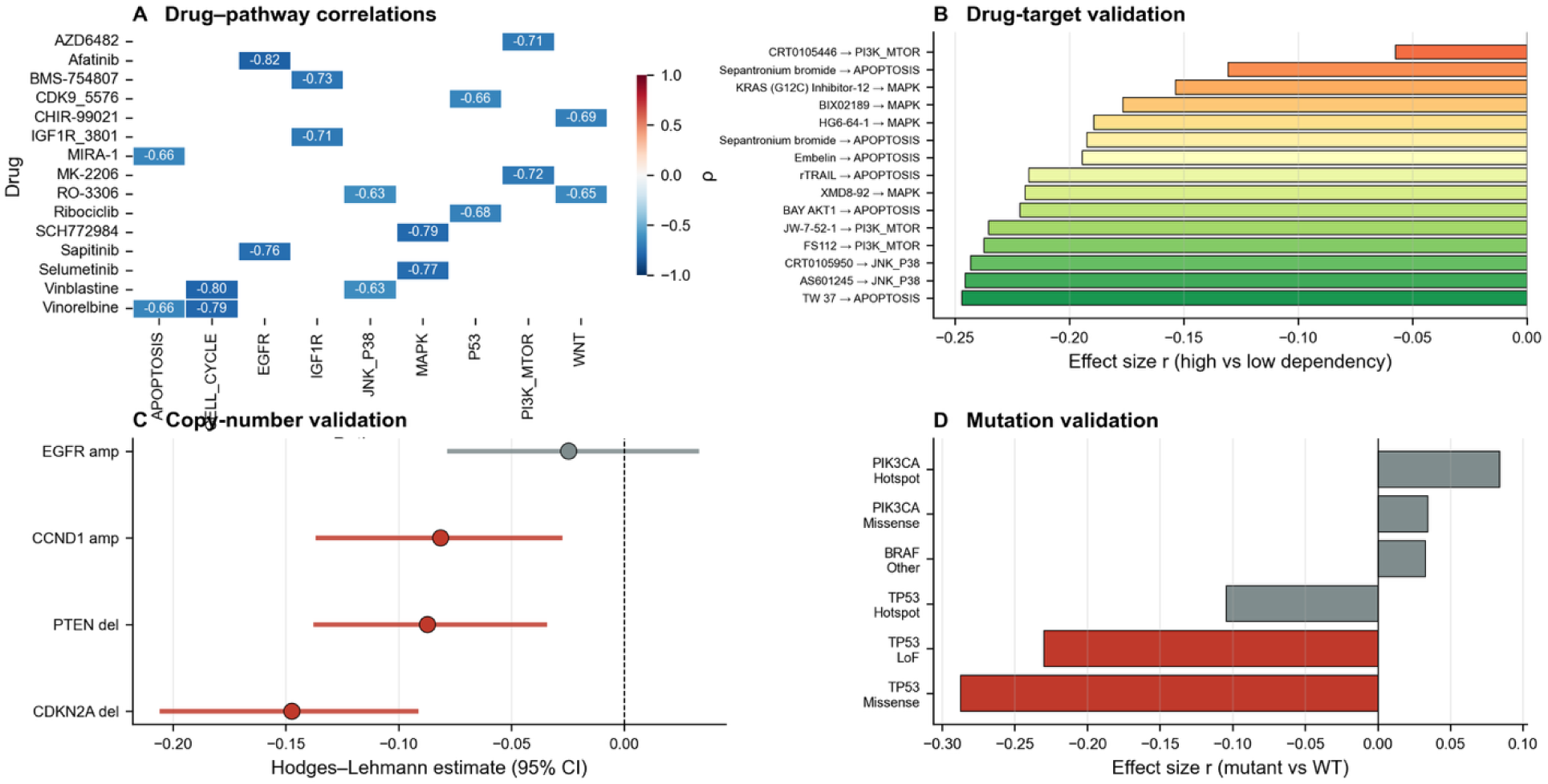
Pharmacological, copy-number, and mutation validation of predicted pathway dependencies. (A) Drug–pathway Spearman correlations. For each drug with ≥10 measured cell lines, the mean LN-IC50 per cell line was correlated with each pathway’s dependency score; for the drug’s own target pathway, dependency was computed leave-one-drug-out to avoid self-correlation. Shown are the two strongest negative correlations per pathway (ρ < −0.3, q < 0.05). The color scale is Spearman ρ. (B) Drug-target validation. For each drug, cell lines were binarised at the 75th percentile of leave-one-drug-out target-pathway dependency (High vs Low). Bars show the effect size r from a one-sided Mann–Whitney U test (alternative = ‘less’, expecting lower IC50 in High-dependency lines); only tests with q < 0.05 are shown. (C) Copy-number validation. For four a priori hypotheses (EGFR amplification → EGFR dependency; PTEN deletion → PI3K/MTOR dependency; CDKN2A deletion → CELL_CYCLE dependency; CCND1 amplification → CELL_CYCLE dependency), predicted dependency was compared between Altered and Normal (diploid) tumors. Points show Hodges–Lehmann estimates; horizontal lines are bootstrap 95% CIs (1,000 resamples). Red points indicate FDR-significant tests (q < 0.05); grey points are non-significant. (D) Mutation validation. Effect sizes (r) from two-sided Mann–Whitney U tests comparing predicted dependency between mutant and wild-type tumors for TP53 → P53, PIK3CA → PI3K/MTOR, and BRAF → MAPK. Only mutation classes with ≥5 samples are shown. Red bars indicate q < 0.05. Abbreviations: CNV, copy-number variation; HL, Hodges–Lehmann; FDR, false-discovery rate

Genomic validation confirmed that predicted dependencies align with DNA-level alterations for three of four tested copy-number events (Table 2, Table S6, Fig. 2C). PTEN deletion (n-alt = 140 vs n-norm= 338), CDKN2A deletion (n-alt = 308 vs n-norm= 145), and CCND1 amplification (n-alt = 224 vs n-norm= 237) were each associated with lower predicted dependency on the matched pathway (PTEN del → PI3K/MTOR, HL = −0.088, r = −0.15, q = 0.001, CDKN2A del → CELL_CYCLE, HL = −0.15, r = −0.23, q = 3× 10⁻^6^; CCND1 amp → CELL_CYCLE HL = −0.08, r = −0.13, q = 0.004). EGFR amplification did not reach significance (HL = −0.024, r = −0.034, q = 0.44). Stratifying mutations by functional type refined these observations (Table S7). In TP53, missense mutations (n = 141, HL = −0.22, r = −0.29, q = 6.13× 10⁻^6^) and loss-of-function mutations (n = 101, HL = −0.18, r = −0.23, q = 2.06×10^−4^) were both associated with lower P53 dependency, whereas TP53 hotspot mutations (n = 54, HL = −0.10, r = −0.10, q = 0.18) trended in the same direction without reaching significance. PIK3CA hotspot mutations (n = 59, HL = +0.09, r = 0.084, q = 0.14) and BRAF non-V600E mutations (n = 8, HL = +0.16, r = 0.03, q = 0.55) did not reach FDR significance. PTEN loss-of-function (n = 6) and missense (n = 5) subsets were small and non-significant. The TP53 result is the most informative: consistent with loss-of-function biology, TP53-mutant tumors show reduced transcriptional dependency on the p53 pathway, which is already disrupted and therefore not a targetable vulnerability (Table 2, Fig. 2D).

**Table 2.**
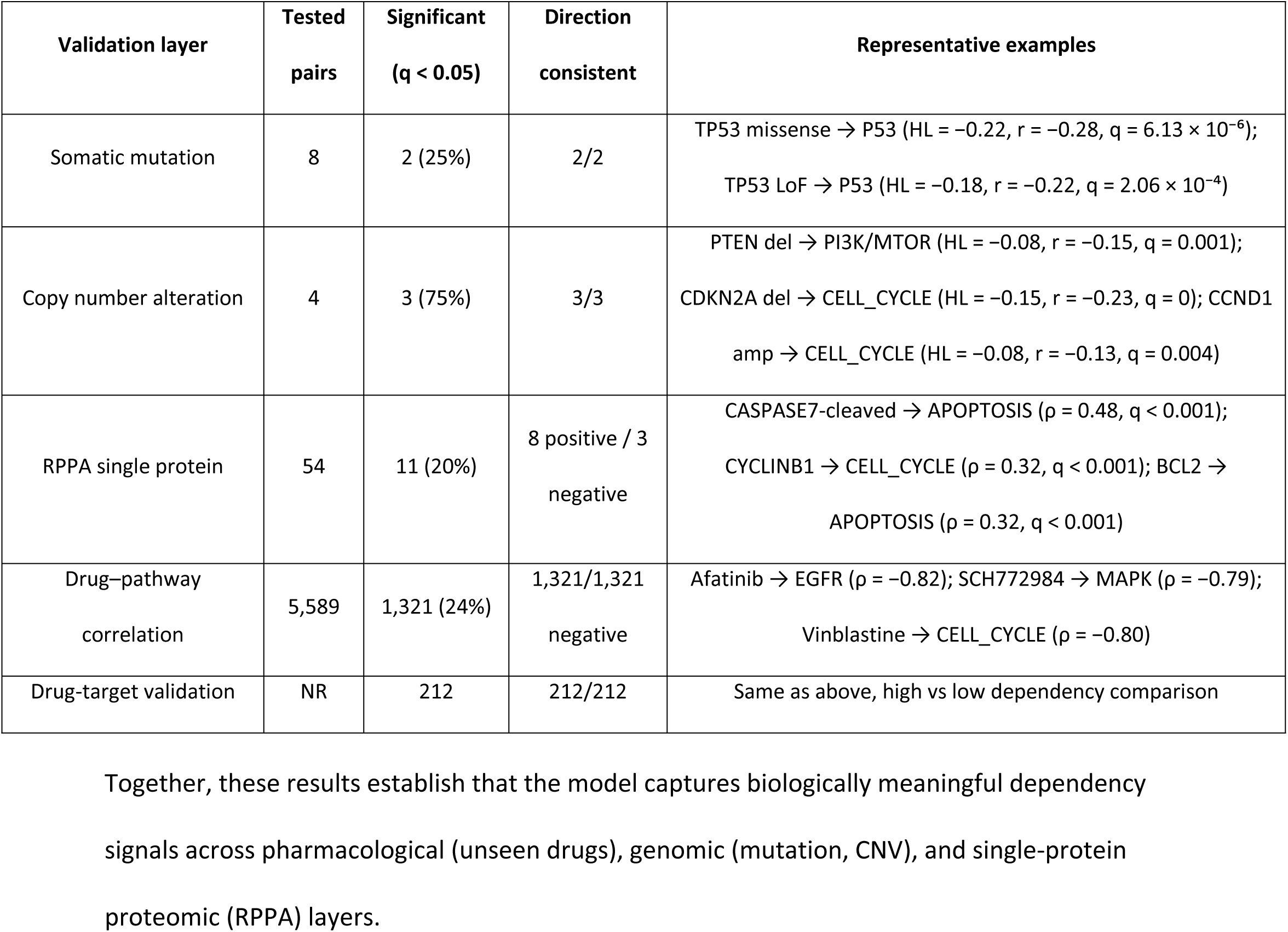
Biological validation of predicted pathway dependencies across pharmacological, genomic, and proteomic layers. Tested pairs, FDR-significant results (q < 0.05), direction consistency, and representative examples for each validation layer.

Proteomic validation involved correlating each predicted dependency score with its matched RPPA protein or phospho-protein at the single-protein level. Eleven pairs were significant after FDR correction (|ρ| = 0.18–0.48, all q < 0.05; Table S8, Table 2, Fig. 3A), including canonical readouts such as cleaved caspase-7 with APOPTOSIS (ρ = 0.48), cyclin B1 with CELL_CYCLE (ρ = 0.32), BCL2 with APOPTOSIS (ρ = 0.32), CRAF with MAPK (ρ = 0.26), HER2 pY1248 with EGFR (ρ = 0.22). Three proteins showed negative correlations with their matched pathway dependency, including BCLX with APOPTOSIS (ρ = −0.18), BAK with APOPTOSIS (ρ = −0.21), and cyclin D1 with CELL_CYCLE (ρ = −0.19).

**Figure 3.**
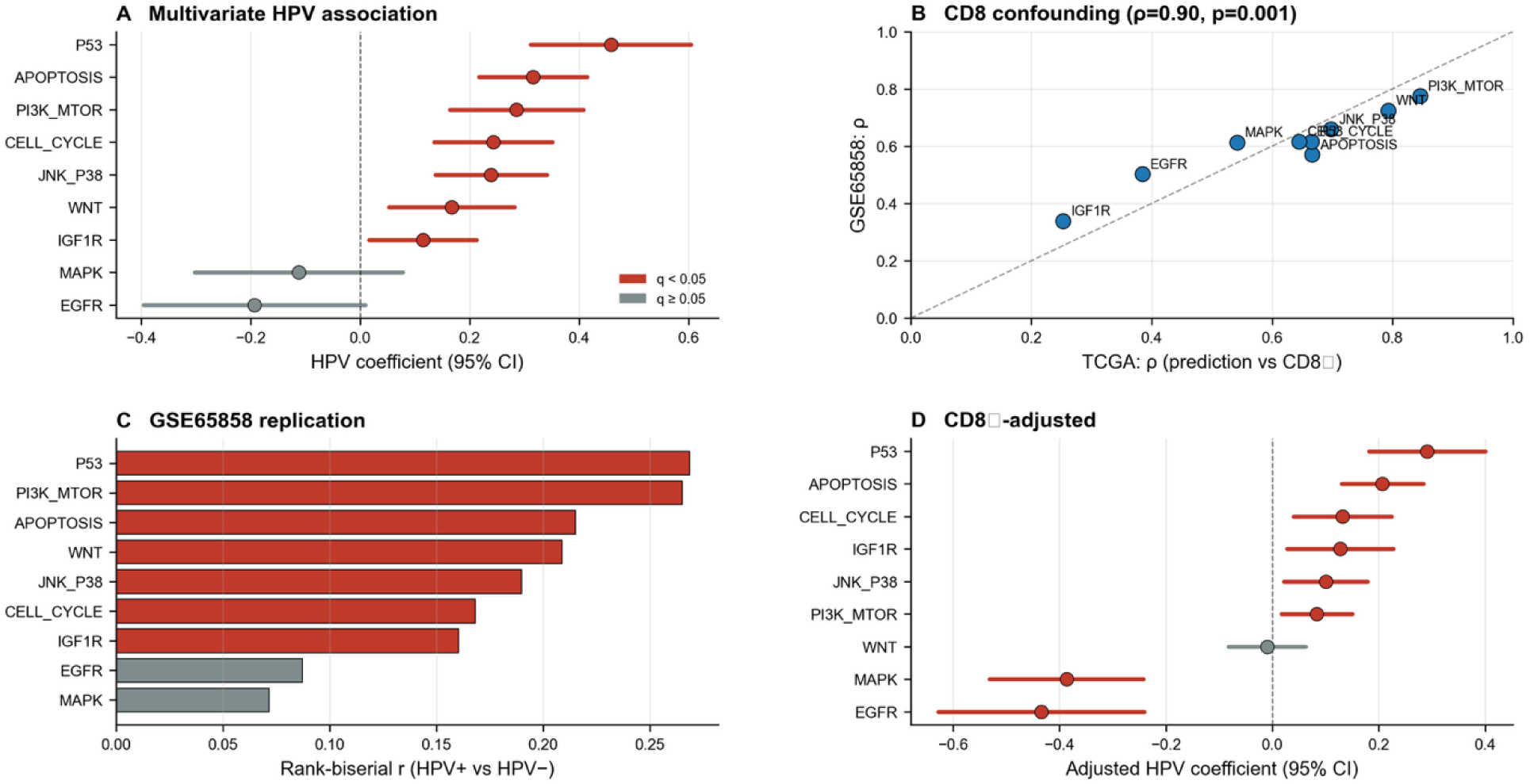
HPV-associated dependency program in TCGA-HNSC and replication in GSE65858. (A) Multivariate HPV association in TCGA-HNSC. For each pathway, an OLS model was fit: dependency ∼ HPV + age + gender + stage + smoking + treatment score (N = 111; 38 HPV+, 73 HPV−; complete cases). Points are HPV coefficients; horizontal lines are 95% CIs. Red points indicate FDR-significant coefficients (q < 0.05). (B) Cross-cohort CD8⁺ confounding. Each point is a pathway. The x-axis is the Spearman ρ between the pathway prediction and the CD8⁺ score in TCGA-HNSC (n = 114); the y- axis is the corresponding ρ in GSE65858 (n = 270). The dashed diagonal indicates perfect agreement; the reported ρ is the cross-cohort Spearman correlation of the confounding magnitudes. (C) Replication of the HPV dependency program in GSE65858 (n = 256; 60 HPV+, 196 HPV−). Bars show the rank-biserial effect size r from a one-sided Mann–Whitney U test (alternative = ‘greater’) comparing HPV+ vs HPV− predictions per pathway. Red bars indicate q < 0.05. (D) CD8⁺-adjusted HPV association in TCGA-HNSC. For each pathway, an OLS model was fit: dependency ∼ HPV + CD8. Points are adjusted HPV coefficients; horizontal lines are 95% CIs. Red points indicate q < 0.05. Abbreviations: HPV, human papillomavirus; OLS, ordinary least squares; FDR, false-discovery rate.

Together, these results establish that the model captures biologically meaningful dependency signals across pharmacological (unseen drugs), genomic (mutation, CNV), and single-protein proteomic (RPPA) layers.

### 2.2 The phenotype dependency program

#### 2.2.1 HPV association in TCGA

In TCGA-HNSC, we compared pathway dependencies between HPV-positive (n = 38) and HPV-negative (n = 73) patients with complete covariate data (n = 111), defining HPV status by p16 immunohistochemistry. To assess independence from clinical covariates, we fitted a multivariate ordinary least squares model for each pathway (dependency ∼ HPV + age + gender + stage + smoking + treatment score) for each pathway (Figure 3A, Table S9). Alcohol consumption was excluded from the primary multivariate model because it was recorded for only 42% of the cohort (240/565), reducing the analyzable sample from 111 to 53 patients. It was instead examined as a phenotype association (Section 3.2.4) and noted as a limitation. Seven of nine pathways showed significantly higher dependency in HPV-positive tumors after FDR correction: P53 (β = 0.46, q = 4.9× 10⁻^8^), APOPTOSIS (β = 0.31, q = 4.9× 10⁻^8^), PI3K/MTOR (β = 0.28, q = 2.35× 10⁻^5^), CELL_CYCLE (β = 0.24, q = 3.63× 10⁻^5^), JNK/P38 (β = 0.24, 2.32× 10⁻^5^), WNT (β = 0.17, q = 7 × 10⁻^3^), and IGF1R (β = 0.11, q = 2.93 × 10⁻^2^). EGFR trended toward lower dependency in HPV-positive tumors (β = −0.19, q = 0.069), and MAPK was not significantly associated with HPV status (β = −0.11, q = 0.24).

To determine which genes within the Hallmark P53 pathway drive the HPV–P53 dependency association, we analyzed the 200-gene Hallmark P53 set. Of these, 190 genes were present in the TCGA expression matrix, including TP53.

#### 2.2.2 Universal CD8+ confounding

Because GDSC cell lines lack immune compartments, training leaves immune-related Hallmark feature weights unconstrained. In bulk tumors, predictions may therefore reflect biological variation in immune features. We tested this by correlating CD8⁺ T-cell infiltration (mean CD8A, CD8B, GZMA, GZMB, and PRF1 expression) with nine pathway predictions in TCGA-HNSC (114 HPV-annotated samples) and GSE65858 (270 samples). Every pathway prediction correlated positively with the CD8⁺ score in both cohorts (Table S10). In TCGA, Spearman ρ ranged from +0.25 (IGF1R) to +0.85 (PI3K/MTOR); in GSE65858, from +0.34 (IGF1R) to +0.78 (PI3K/MTOR).

All correlations were significant after FDR correction (q < 10⁻⁸). The ranking of pathways by confounding magnitude was nearly identical across cohorts (Spearman ρ = +0.90, p = 0.0009), with PI3K/MTOR and WNT at the top in both, and IGF1R at the bottom in both. This cross-cohort reproducibility indicates that the confounding is a property of the model and its application to bulk tumor transcriptomes, not a cohort- or platform-specific artifact. The magnitude of confounding varied by pathway and is relevant to the interpretation of downstream analyses: PI3K/MTOR and WNT predictions were heavily dominated by immune infiltration in patient tumors (ρ > 0.72 in both cohorts), whereas IGF1R was only weakly affected (ρ < 0.34 in both). Intermediate confounding was observed for JNK/P38, P53, APOPTOSIS, CELL_CYCLE, MAPK, and EGFR (ρ = 0.38–0.70) (Fig.3B).

#### 2.2.3 External replication in GSE65858

To test whether the HPV dependency program extends beyond TCGA, we applied the same models and hyperparameters to GSE65858 (n = 256; 60 HPV-positive, 196 HPV-negative) and repeated univariate HPV association analysis with FDR correction (Table 3, Table S11). GSE65858 covariates were not adjusted, preserving a common specification with the confounder-adjusted TCGA model and enabling direct comparison of effect directions. All seven pathways significantly associated with HPV status in TCGA replicated in GSE65858 with concordant direction of effect: P53 (r = 0.27, q < 0.001), PI3K/MTOR (r = 0.26, q < 0.001), APOPTOSIS (r = 0.21, q = 0.002), WNT (r = 0.21, q = 0.002), JNK/P38 (r = 0.19, q = 0.004), CELL_CYCLE (r = 0.17, q = 0.011), and IGF1R (r = 0.16, q = 0.013). EGFR (r = 0.09, q = 0.18) and MAPK (r = 0.07, q = 0.25) did not reach significance in either cohort. Effect sizes were modest (r = 0.16–0.27), but the direction of effect was concordant across all seven replicated pathways, establishing that the HPV dependency program generalizes beyond TCGA (Fig.3C).

**Table 3.**
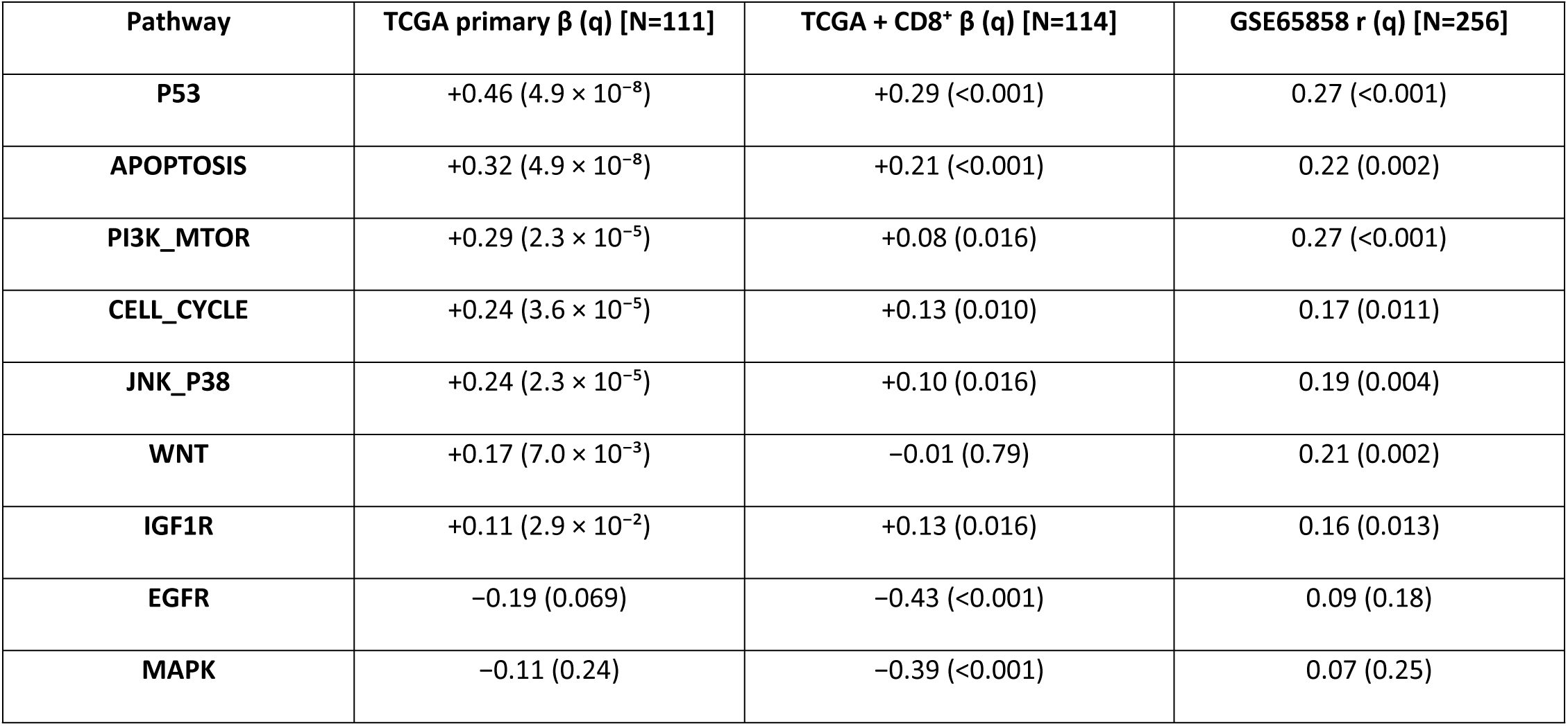
HPV-positive vs HPV-negative pathway dependencies in TCGA-HNSC and GSE65858. TCGA primary: OLS β adjusted for age, gender, stage, smoking, and treatment (N = 111). TCGA + CD8⁺: OLS β adjusted for CD8⁺ only (N = 114). GSE65858: one-sided Mann–Whitney effect size r (N = 256). All q-values BH-FDR-corrected across nine pathways.

#### 2.2.4 CD8-adjusted HPV association

Because CD8⁺ infiltration confounded pathway predictions to varying degrees (Section 2.2.2), we modeled each pathway as dependency ∼ HPV + CD8 to isolate the immune-independent HPV association. The CD8⁺-adjusted coefficient β_HPV estimates HPV’s effect on dependency at fixed CD8⁺ infiltration. This model included only HPV and CD8; the primary analysis adjusted for age, gender, stage, smoking, and treatment score) is reported separately in Section 2.2.1. After CD8⁺ adjustment, eight of nine pathways remained significantly associated with HPV status (only WNT became null; q = 0.79). The retained β_HPV was strongly positive for P53 (β = +0.30, q < 0.001) and APOPTOSIS (β = +0.20, q < 0.001); moderately positive for CELL_CYCLE (β = +0.13, q = 5.28 × 10⁻^3^) and IGF1R (β = +0.13, q = 0.016); and weakly positive for JNK/P38 (β = +0.10, q = 0.015) and PI3K/MTOR (β = +0.08, q = 0.015). Two RTK-driven pathways shifted to significantly negative coefficients: EGFR (β = −0.43, q < 0.001) and MAPK (β = −0.39, q < 0.001). Retention tracked CD8⁺ confounding: the most confounded pathways, PI3K/MTOR and WNT, attenuated most after adjustment (28% of the unadjusted effect remained for PI3K/MTOR; WNT became null). The least confounded, IGF1R, was unchanged (β = +0.11 unadjusted, +0.13 adjusted). P53 and APOPTOSIS (ρ ≈ 0.67) retained 63% and 66% of their effects. This gradient is expected when adjustment isolates each HPV association’s immune-independent component; the residual fraction reflects its intrinsic association with immune infiltration (Table 3, Table S12, Fig.3D).

#### 2.2.5 Smoking sensitivity for EGFR/MAPK

The EGFR and MAPK sign reversal after CD8⁺ adjustment may reflect smoking confounding, as smoking is enriched in HPV-negative tumors and activates RTK signaling. We tested this by adding smoking to the CD8⁺-adjusted model in the same sample (N = 112; two patients excluded for missing smoking status). The HPV coefficients were unchanged (EGFR: β = −0.446 → −0.446; MAPK: β = −0.391 → −0.383), with no change in effect size or FDR significance (both q < 0.001). The suppression effect is therefore not driven by smoking confounding (Table S13).

#### 2.2.6 Non-HPV clinical associations

To distinguish HPV-driven dependencies from those associated with other variables, we tested stage, treatment, age, smoking, and alcohol against nine pathway dependencies (Fig. S4A & S4B).

No pathway was associated with clinical stage after FDR correction. CELL_CYCLE showed the strongest nominal trend (Spearman ρ = 0.10, unadjusted p = 0.01), with higher predicted dependency in higher-stage tumors, but this was not significant after correction (q = 0.33). Patients were grouped by treatment intensity: high (≥ 2 modalities), low (1), or none (0). No pathway differed between high- and low-intensity groups after FDR correction (all q > 0.80; Table S15). Across 36 tests (nine pathways × four covariates), no age, smoking, or alcohol association survived correction (all q > 0.05; Table S14); smoking and alcohol were unassociated with every pathway (all |ρ| < 0.11). These findings indicate that HPV status, rather than classical clinical covariates, primarily drives the dependency program.

#### 2.2.7 Resolution of the P53 paradox: evidence for residual p53 activity in HPV-positive tumors

HPV-positive tumors show greater predicted P53 pathway dependency (Section 2.2.1), despite HPV E6’s known targeting of p53 for ubiquitin-mediated degradation. Because the model predicts dependency on the 190-gene Hallmark P53 transcriptional program rather than on p53 abundance, we examined which components drive this association.

Across 190 Hallmark P53 genes in the TCGA matrix (N = 114), we fit P53_dependency ∼ HPV + gene + Age + Gender. No gene eliminated the HPV association after FDR correction (all q_HPV_adj < 0.05). Compared with 500 same-size, expression-matched random sets, Hallmark P53 genes showed greater median attenuation (0.040 vs. 0.019; empirical p < 0.0001), supporting program-level rather than single-gene specificity. The top attenuators were predominantly p53 transcriptional targets upregulated in HPV-positive tumors (POLH, TP53, ISCU, XPC, CYFIP2, TSPYL2) and p53-repressed genes downregulated in HPV-positive tumors (SLC7A11, SLC3A2, UPP1, SDC1, TGFA, HBEGF, KLK8, KRT17) (Fig.4A, Fig.S8A, Table S16).

**Figure 4.**
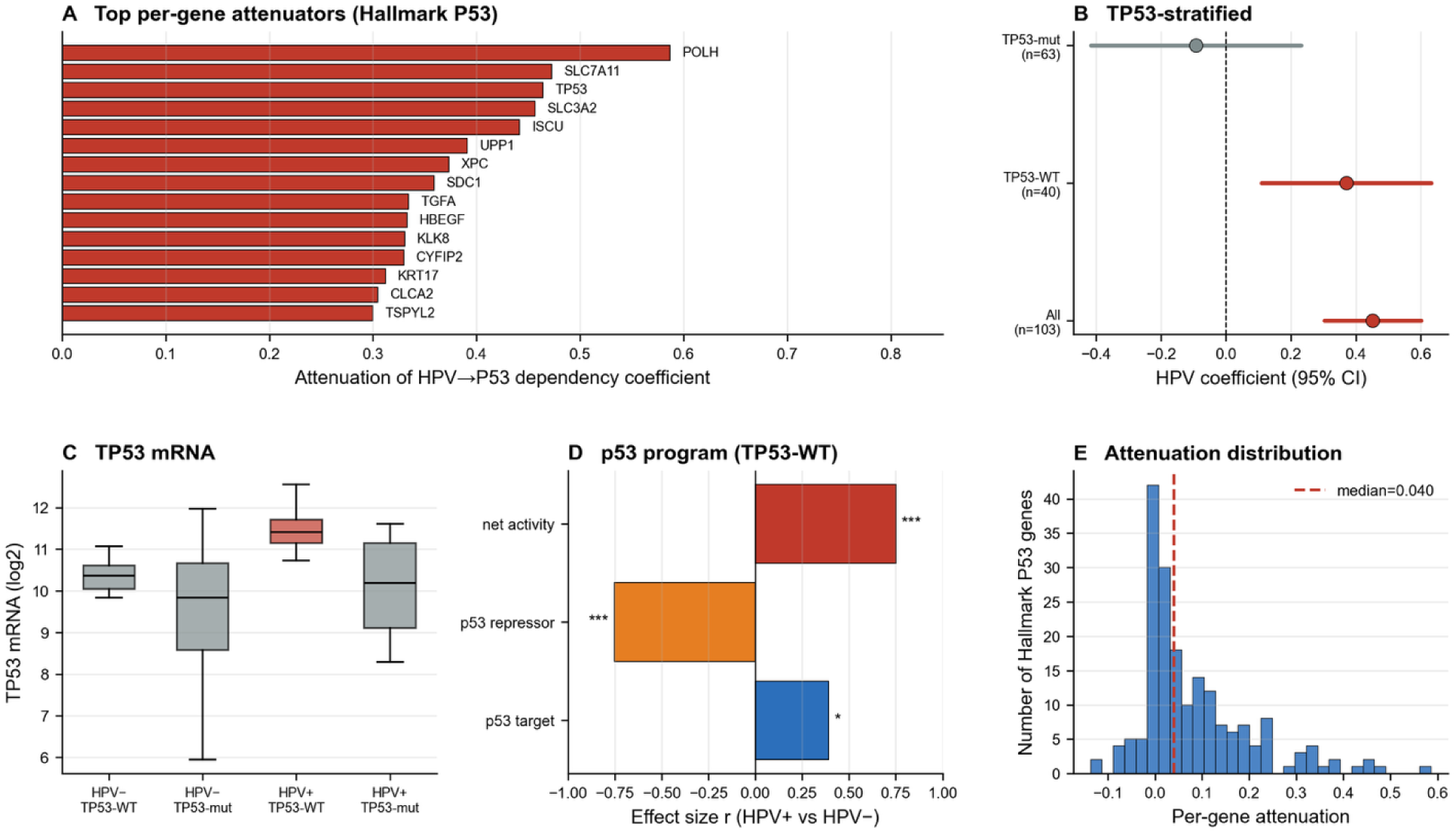
Resolution of the P53 paradox: evidence for residual p53 transcriptional activity in HPV-positive HNSCC. (A) Per-gene mediation of the HPV → P53 dependency association across the 190 Hallmark P53 genes present in TCGA-HNSC. Bars show the attenuation of the HPV coefficient when each gene is added to the model (attenuation = (β_base − β_adj) / β_base). Red bars indicate genes with FDR-significant gene effects (q_gene < 0.05). The 15 genes with the largest attenuation are labeled. (B) TP53-stratified HPV association with P53 dependency. Points are HPV coefficients from OLS models (P53_dependency ∼ HPV + age + gender) within All (n = 114), TP53-WT (n = 40), and TP53-mut (n = 4) strata; horizontal lines are 95% CIs. Red points indicate p < 0.05. (C) TP53 mRNA by HPV and TP53 mutation status. Boxes show median and interquartile range; whiskers extend to 1.5 × IQR. The red box highlights HPV+/TP53-WT tumors. (D) p53 transcriptional program scores in TP53-WT tumors. Bars show effect sizes (r) from Mann–Whitney U tests comparing HPV+ vs HPV− tumors for the p53 target score (13 canonical p53 targets), p53 repressor score (12 p53-repressed genes), and net activity score (targets − repressors). Asterisks indicate FDR-corrected significance (q < 0.05). (E) Distribution of per-gene attenuation across the 190 Hallmark P53 genes. The dashed red line marks the median. Abbreviations: HPV, human papillomavirus; WT, wild-type; FDR, false-discovery rate.

We formalized this observation by computing a p53 target score (mean of 13 canonical p53 targets) and a p53 repressor score (mean of 12 p53-repressed genes). Within TP53-WT tumors, HPV-positive tumors showed higher p53 target expression (r = 0.39, q = 0.014) and lower p53-repressed gene expression (r = −0.75, q < 0.001), yielding a net p53 activity score that was 2.5 log units higher in HPV-positive than HPV-negative tumors (r = 0.75, q < 0.001). This signature was stronger in the TP53-WT stratum than in the full cohort, indicating that the TP53-mutant minority does not inflate the association (Fig.4D, Table S17).

To isolate the driver of the association, we stratified by TP53 mutation status. Within TP53-WT tumors (n = 40), HPV remained significantly associated with higher P53 dependency (β = +0.37, 95% CI 0.11–0.63, p = 0.007, q = 0.010). The TP53-mutant stratum contained only four HPV-positive tumors and was uninformative. The TP53-WT result establishes that the HPV association is not driven by the mutant minority (Table S18).

TP53 mRNA was highest in HPV-positive/TP53-WT tumors (median 11.41 vs. 10.36 in HPV-negative/TP53-WT, p < 0.0001). Within HPV-positive/TP53-WT tumors (n = 28), TP53 mRNA correlated with the p53 target score (Spearman ρ = 0.417, 95% CI 0.05–0.68, ρ² = 0.17, p = 0.027), consistent with a model in which E6-mediated p53 protein degradation is partially compensated by p53 gene expression upregulation. This correlation is bidirectional by design (TP53 is itself a p53 target), so it is consistent with the compensatory mechanism but does not establish it (Fig.4C).

### 2.3 Pathway co-dependency architecture

#### 2.3.1 Co-dependency network in the full cohort

Across the full TCGA-HNSC cohort (n = 565), 30 of 36 pathway pairs exhibited significant positive correlations after FDR correction (|ρ| > 0.3, q < 0.05; Table 4). The most strongly coupled pairs were CELL_CYCLE↔JNK_P38 (ρ = 0.94), PI3K_MTOR↔WNT (ρ = 0.92), JNK_P38↔WNT (ρ = 0.88), APOPTOSIS↔CELL_CYCLE (ρ = 0.86), and JNK_P38↔PI3K_MTOR (ρ = 0.84). A second tier of strong coupling included CELL_CYCLE↔WNT (ρ = 0.83), APOPTOSIS↔JNK_P38 (ρ = 0.81), CELL_CYCLE↔PI3K_MTOR (ρ = 0.80), APOPTOSIS↔P53 (ρ = 0.78), and APOPTOSIS↔PI3K_MTOR (ρ = 0.78) (Figure 5A, Table S19). Only one pathway pair showed a significant negative correlation: EGFR↔IGF1R (ρ = −0.15, q = 3.2 × 10⁻⁴).

**Figure 5.**
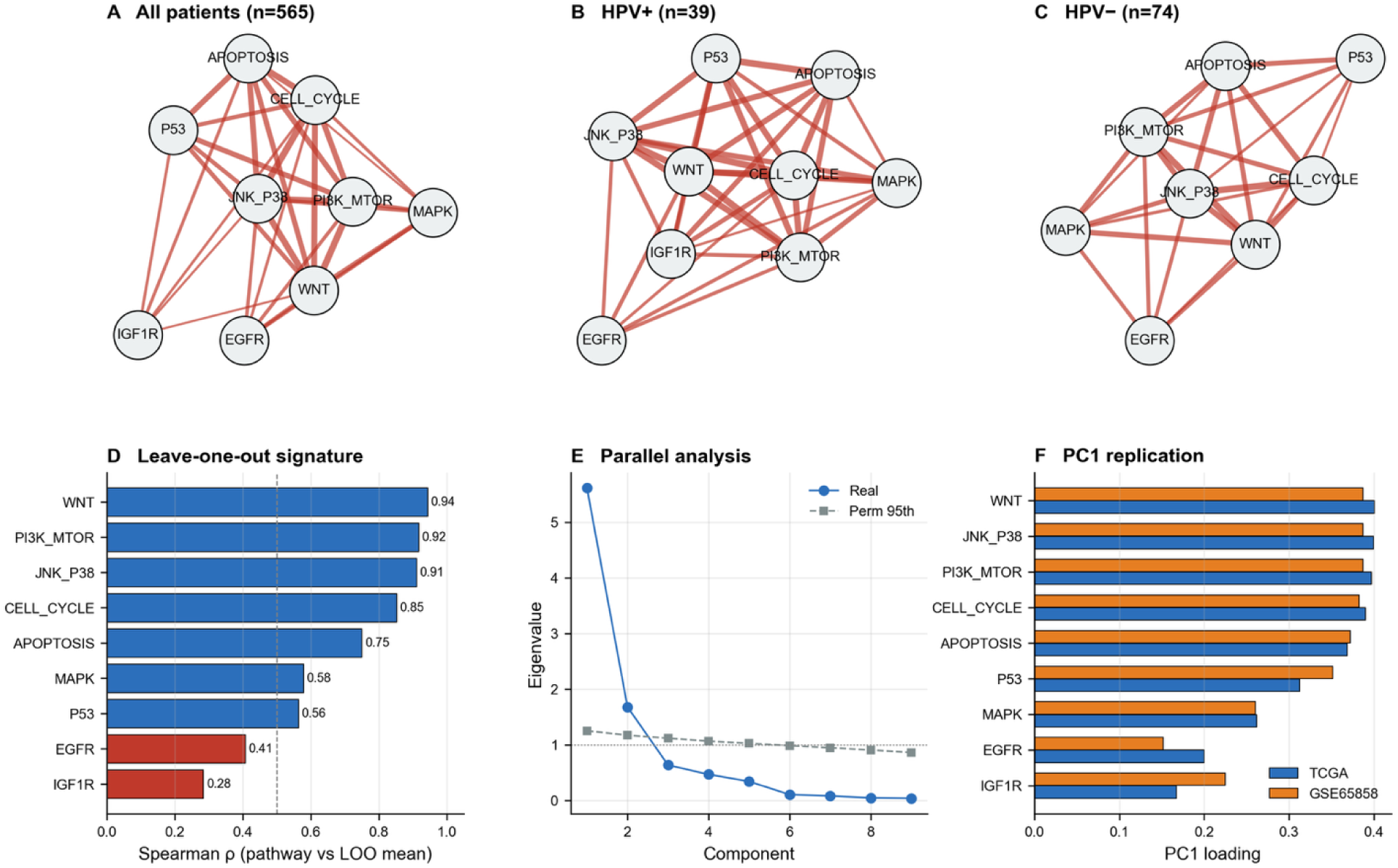
Co-dependency architecture, global dependency axis, and replication in GSE65858. (A–C) Pathway co-dependency networks in TCGA-HNSC. Nodes are the nine predicted pathway dependencies; edges are significant pairwise Spearman correlations (|ρ| > 0.3, q < 0.05, FDR-corrected). Edge width is proportional to |ρ|; red edges indicate positive correlations, blue edges negative. Networks are shown for (A) the full cohort (n = 565), (B) HPV-positive tumors (n = 39), and (C) HPV-negative tumors (n = 74). (D) Leave-one-out (LOO) global dependency signature. For each pathway, the LOO score was computed as the mean of the other eight z-scored pathway predictions; bars show the Spearman ρ between each pathway and its LOO score. All correlations were significant after FDR correction. (E) Parallel analysis of the nine-pathway dependency matrix (TCGA-HNSC, n = 565). Solid line: observed eigenvalues; dashed line: 95th percentile of a permutation null (1,000 permutations, each pathway independently shuffled); dotted line: permuted median; horizontal dotted line: λ = 1. (F) Replication of PC1 loadings in GSE65858. Bars compare PC1 loadings from TCGA-HNSC (blue) and GSE65858 (orange) for each pathway. Tucker congruence between the two loading vectors is reported in the main text. Abbreviations: FDR, false-discovery rate; LOO, leave-one-out; PC1, first principal component.

**Table 4.** Pairwise Spearman correlations between predicted pathway dependencies in TCGA-HNSC. Only pairs significant in the clinically annotated cohort (|ρ| > 0.3, q < 0.05) are shown (N = 113; 39 HPV+, 74 HPV−). Replication indicates whether the pair also met criteria in the corresponding GSE65858 subgroup.

| Pathway A | Pathway B | $\rho$ | 95% CI | $q$ | Replicated HPV+ | Replicated HPV- |
| --- | --- | --- | --- | --- | --- | --- |
| CELL_CYCLE | JNK_P38 | 0.937 | 0.926 – 0.946 | 1.53e-257 | Yes (0.935) | Yes (0.907) |
| PI3K_MTOR | WNT | 0.922 | 0.909 – 0.934 | 4.89e-233 | Yes (0.954) | Yes (0.918) |
| JNK_P38 | WNT | 0.881 | 0.861 – 0.898 | 5.94e-184 | Yes (0.940) | Yes (0.817) |
| APOPTOSIS | CELL_CYCLE | 0.86 | 0.836 – 0.880 | 2.37e-165 | Yes (0.869) | Yes (0.797) |
| JNK_P38 | PI3K_MTOR | 0.841 | 0.816 – 0.864 | 9.62e-152 | Yes (0.878) | Yes (0.784) |
| CELL_CYCLE | WNT | 0.826 | 0.798 – 0.850 | 1.86e-141 | Yes (0.906) | Yes (0.709) |
| APOPTOSIS | JNK_P38 | 0.808 | 0.778 – 0.835 | 5.89e-131 | Yes (0.812) | Yes (0.768) |
| CELL_CYCLE | PI3K_MTOR | 0.799 | 0.767 – 0.827 | 9.16e-126 | Yes (0.877) | Yes (0.694) |
| APOPTOSIS | P53 | 0.784 | 0.750 – 0.814 | 3.07e-118 | Yes (0.939) | Yes (0.705) |
| APOPTOSIS | PI3K_MTOR | 0.781 | 0.747 – 0.811 | 1.20e-116 | Yes (0.804) | Yes (0.756) |
| P53 | PI3K_MTOR | 0.748 | 0.709 – 0.782 | 7.71e-102 | Yes (0.870) | Yes (0.607) |
| APOPTOSIS | WNT | 0.733 | 0.692 – 0.769 | 9.78e-96 | Yes (0.798) | Yes (0.685) |
| MAPK | WNT | 0.676 | 0.629 – 0.719 | 2.18e-76 | Yes (0.807) | Yes (0.770) |
| P53 | WNT | 0.62 | 0.566 – 0.668 | 8.31e-61 | Yes (0.831) | Yes (0.492) |
| CELL_CYCLE | P53 | 0.595 | 0.539 – 0.646 | 5.38e-55 | Yes (0.855) | Yes (0.350) |
| JNK_P38 | P53 | 0.593 | 0.537 – 0.644 | 1.15e-54 | Yes (0.826) | Yes (0.397) |
| JNK_P38 | MAPK | 0.592 | 0.535 – 0.643 | 2.31e-54 | Yes (0.754) | Yes (0.658) |
| EGFR | WNT | 0.559 | 0.500 – 0.613 | 1.74e-47 | Yes (0.564) | Yes (0.600) |
| MAPK | PI3K_MTOR | 0.551 | 0.491 – 0.606 | 6.68e-46 | Yes (0.700) | Yes (0.667) |
| EGFR | MAPK | 0.541 | 0.480 – 0.597 | 4.80e-44 | Yes (0.512) | Yes (0.543) |
| EGFR | PI3K_MTOR | 0.503 | 0.439 – 0.562 | 2.57e-37 | Yes (0.511) | Yes (0.559) |
| EGFR | JNK_P38 | 0.476 | 0.410 – 0.537 | 4.40e-33 | Yes (0.490) | Yes (0.521) |
| CELL_CYCLE | MAPK | 0.453 | 0.385 – 0.516 | 9.48e-30 | Yes (0.638) | Yes (0.447) |
| APOPTOSIS | IGF1R | 0.452 | 0.384 – 0.515 | 1.32e-29 | Yes (0.720) | — |
| IGF1R | P53 | 0.404 | 0.332 – 0.471 | 2.07e-23 | Yes (0.678) | — |
| CELL_CYCLE | EGFR | 0.401 | 0.330 – 0.468 | 4.04e-23 | Yes (0.422) | Yes (0.395) |
| CELL_CYCLE | IGF1R | 0.38 | 0.307 – 0.448 | 1.01e-20 | Yes (0.648) | — |
| IGF1R | JNK_P38 | 0.33 | 0.255 – 0.402 | 1.02e-15 | Yes (0.571) | — |
| IGF1R | WNT | 0.308 | 0.231 – 0.381 | 8.77e-14 | Yes (0.529) | — |
| APOPTOSIS | MAPK | 0.303 | 0.226 – 0.376 | 2.26e-13 | Yes (0.454) | Yes (0.388) |

**Table 5.** Pairwise pathway co-dependency before and after removal of the global dependency axis (PC1). Spearman correlations were calculated for z-scored pathway dependency predictions in TCGA-HNSCC (n = 565). ρ_resid is the correlation after regressing each pathway on PC1. Δ|ρ| (%) = (|ρ_full| − |ρ_resid|) / |ρ_full| × 100; negative values indicate increased absolute correlation. “Sign flip” means ρ_full and ρ_resid have opposite signs. Pairs are sorted by ρ_full. Bold values (if used in the final typeset table) mark pairs with |ρ_resid| > 0.3.

| Pathway pair | $\rho_{full}$ | $\rho_{resid}$ | $\Delta\vert \rho\vert $ (%) | Sign flip |
| --- | --- | --- | --- | --- |
| CELL_CYCLE ↔ JNK_P38 | 0.94 | 0.54 | 42.5 | No |
| PI3K_MTOR ↔ WNT | 0.92 | 0.22 | 75.8 | No |
| JNK_P38 ↔ WNT | 0.88 | -0.13 | 84.7 | Yes |
| APOPTOSIS ↔ CELL_CYCLE | 0.86 | 0.32 | 63.0 | No |
| JNK_P38 ↔ PI3K_MTOR | 0.84 | -0.49 | 41.8 | Yes |
| CELL_CYCLE ↔ WNT | 0.83 | -0.34 | 58.3 | Yes |
| APOPTOSIS ↔ JNK_P38 | 0.81 | -0.08 | 90.7 | Yes |
| CELL_CYCLE ↔ PI3K_MTOR | 0.80 | -0.57 | 28.4 | Yes |
| APOPTOSIS ↔ P53 | 0.78 | 0.42 | 46.1 | No |
| APOPTOSIS ↔ PI3K_MTOR | 0.78 | -0.2 | 73.9 | Yes |

#### 2.3.2 Subgroup analyses

In HPV-positive tumors (n = 39), 33 of 36 pairs were significant, with the strongest coupling between PI3K_MTOR and WNT (ρ = 0.95), JNK_P38 and WNT (ρ = 0.94), APOPTOSIS and P53 (ρ = 0.94), and CELL_CYCLE and JNK_P38 (ρ = 0.94) (Fig. 5B). In HPV-negative tumors (n = 74), 25 of 36 pairs were significant, with the same core module but uniformly weaker coupling (PI3K_MTOR↔WNT ρ = 0.92; CELL_CYCLE↔JNK_P38 ρ = 0.91) (Fig.5C). Stratifying by stage showed 15 of 36 significant pairs in early-stage tumors (n = 21) and 29 of 36 in late-stage tumors (n = 92) (Fig. S5A, Fig. S5B).

#### 2.3.3 Global dependency signature

To quantify the co-dependency network’s dominant axis, we calculated each patient’s leave-one-out (LOO) global dependency score (mean of the other eight pathway predictions) and correlated each pathway with it (Fig. 5D). WNT (ρ = 0.94), PI3K/MTOR (ρ = 0.92), and JNK/P38 (ρ = 0.91) correlated most strongly, followed by CELL_CYCLE (ρ = 0.85), APOPTOSIS (ρ = 0.75), MAPK (ρ = 0.58), P53 (ρ = 0.56), EGFR (ρ = 0.41), and IGF1R (ρ = 0.28); all correlations were significant after FDR correction.

PCA of z-scored TCGA pathway predictions (n = 565) showed that PC1 explained 62.3% of variance, with all nine loadings positive (WNT 0.400, JNK/P38 0.400, PI3K/MTOR 0.397, CELL_CYCLE 0.390, APOPTOSIS 0.368, P53 0.313, MAPK 0.262, EGFR 0.200, IGF1R 0.167; Fig. 5B). This exceeded the permutation null (median PC1 variance 13.3%; 95th percentile 13.9%), and bootstrap analysis (1,000 resamples) placed every loading’s 95% CI above zero. Internal consistency was high (Cronbach’s α = 0.911, 95% CI 0.900–0.922). The signature replicated in GSE65858 (n = 270): PC1 explained 66.7% of variance, with a nearly identical loading vector (Tucker congruence = 0.996; Fig. 5F).

Because the nine pathway predictions correlate with CD8⁺ T-cell infiltration to varying degrees (Section 2.2.2), we tested whether immune infiltration drove the dominant axis. PC1 correlated with the CD8⁺ score (ρ = +0.80, p < 10⁻¹²⁸, n = 565). After regressing each prediction on CD8⁺ and rerunning PCA on residuals, PC1 variance fell from 62.3% to 48.2%, and the residual axis could not be stably estimated across bootstrap resamples. Thus, the dominant axis likely represents a shared transcriptional signal dominated by immune infiltration, not a pathway-intrinsic co-dependency program. Since both cohorts show CD8⁺ infiltration, PC1’s cross-cohort reproducibility reflects this shared microenvironment signal.

We tested whether the weaker second-order structure formed stable modules. A two-factor EFA (varimax; 75.3% cumulative variance) grouped APOPTOSIS, CELL_CYCLE, JNK/P38, PI3K/MTOR, P53, and WNT in a large module, and EGFR and MAPK in a smaller one; IGF1R loaded on neither. Split-half replication confirmed stability (mean Tucker congruence = 0.993 ± 0.030). However, the second factor explained only ∼15% of reliable variance (bifactor ω_subscale = 0.150), and four methods (CFA, EFA, EGA on partial correlations, and Louvain detection on raw correlations) yielded different groupings. Only CELL_CYCLE↔JNK/P38 co-clustered across all methods. Thus, the dependency architecture is not method-independently modular (Fig. 5E).

Because the nine pathway scores co-vary strongly, they are unsuitable as independent predictors: regressing each on the other eight (OLS) yielded a median R² of 0.885 (range: 0.332 for IGF1R to 0.930 for CELL_CYCLE) and mean VIFs of 6.5–9.2. We therefore used a single composite global dependency score. In TCGA-HNSCC, pathway dependencies are dominated by a reproducible, largely immune-infiltration axis; weak residual couplings do not form method-independent modules (Table S19).

#### 2.3.4 Co-occurrence replication in GSE65858

To test whether the co-dependency structure in TCGA-HNSCC replicates in an independent cohort, we computed pairwise Spearman correlations among the nine pathway dependencies in GSE65858, stratified by HPV status, and compared them to TCGA structures (Table S20). A pair was considered replicated if the sign matched TCGA, |ρ| > 0.2, and q < 0.05.

Under the z-scored target, replication was complete in both strata: all 33 TCGA-significant pairs replicated in HPV-positive tumors (n = 60), and all 25 replicated in HPV-negative tumors (n = 196). No TCGA-significant pair failed to replicate. The strongest agreements in HPV+: PI3K_MTOR↔WNT (TCGA ρ = 0.954, GSE ρ = 0.920), JNK_P38↔WNT (TCGA ρ = 0.940, GSE ρ = 0.933), and CELL_CYCLE↔JNK_P38 (TCGA ρ = 0.935, GSE ρ = 0.952). In HPV−: PI3K_MTOR↔WNT (TCGA ρ = 0.918, GSE ρ = 0.882), CELL_CYCLE↔JNK_P38 (TCGA ρ = 0.907, GSE ρ = 0.937), and JNK_P38↔WNT (TCGA ρ = 0.817, GSE ρ = 0.873). Because the shared microenvironment signal is present in both cohorts, this near-complete replication is expected and confirms that the confounding structure is reproducible and platform-independent.

#### 2.3.5 Proteomic co-activity validation

Predicted co-dependency and RPPA directional co-activity showed no cross-layer agreement at the pair level (Spearman ρ across 36 pairs = −0.244, p = 0.36; Figure 6B, Table S21). PCA of nine RPPA directional pathway scores (n = 212) showed that PC1 explained 25.2% of variance (predicted PC1: 61.3%), exceeding the permutation null (95th percentile: 15.8%) but falling well short of prediction. The axes were partly aligned (Tucker congruence = 0.488) yet structurally distinct: predicted PC1 was positive across all pathways, while RPPA PC1 was bipolar, with negative IGF1R (−0.439) and MAPK (−0.282) loadings opposed to positive PI3K/MTOR (0.414), CELL_CYCLE (0.435), EGFR (0.417), and APOPTOSIS (0.369) loadings. Since the predicted co-dependency axis is largely immune-driven (Section 2.3.3), whereas RPPA measures protein abundance and phosphorylation, this cross-layer mismatch supports the interpretation that the prediction captures a shared microenvironment signal, not protein-level pathway co-regulation.

**Figure 6.**
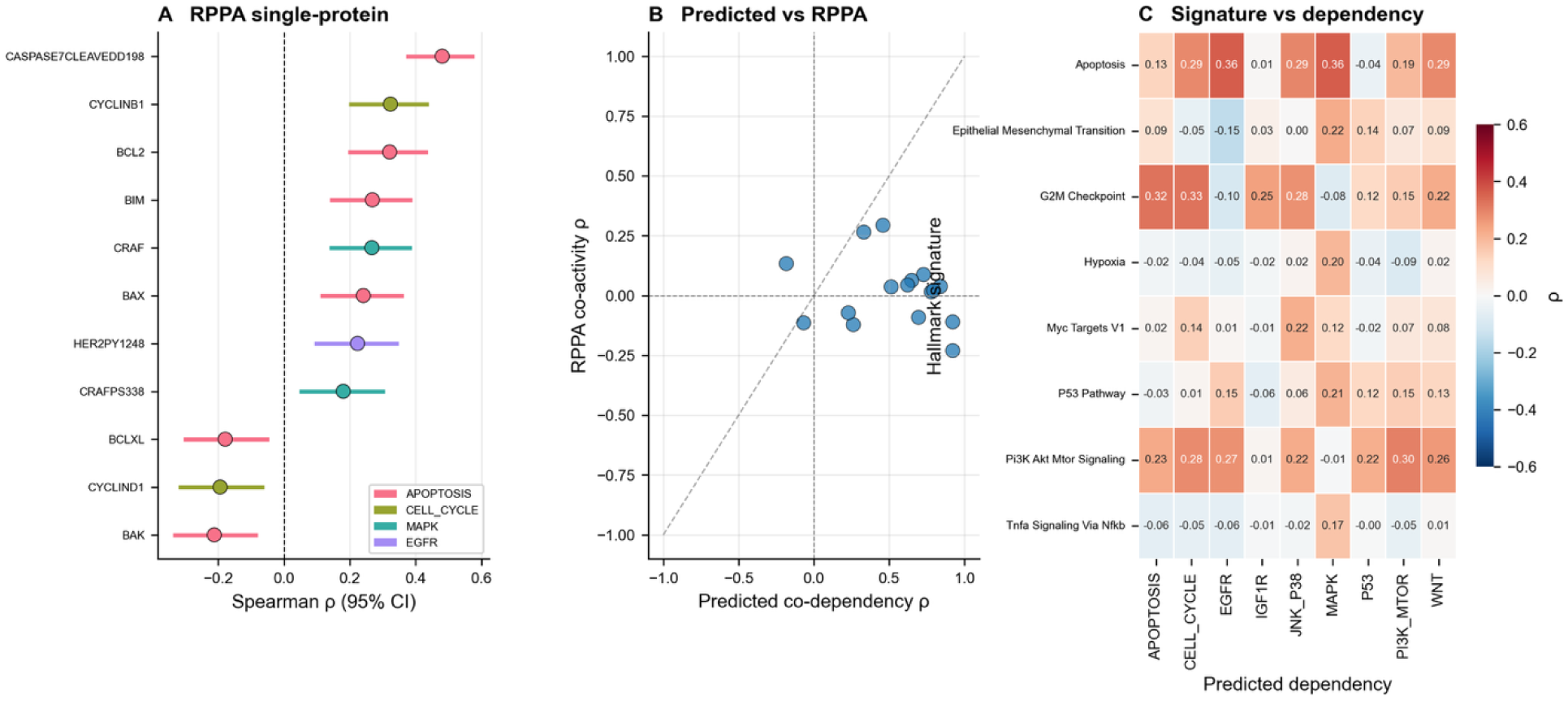
Proteomic validation and cross-layer concordance of predicted pathway dependencies. (A) RPPA single-protein validation. For each protein or phospho-protein mapped to a pathway, Spearman ρ between protein abundance and the matched pathway’s predicted dependency is shown. Only pairs significant after FDR correction (q < 0.05) are plotted. Points are coloured by pathway; horizontal lines are Fisher Z 95% CIs. (B) Cross-layer comparison of pairwise pathway co-activity. Each point is a pathway pair (36 pairs total). The x-axis is the predicted co-dependency ρ (TCGA-HNSC, n = 212 with matched RPPA); the y-axis is the RPPA directional co-activity ρ. The dashed diagonal indicates perfect agreement. (C) Hallmark signature vs predicted dependency heatmap. Spearman correlations between eight selected Hallmark ssGSEA signature scores and the nine predicted pathway dependencies in TCGA-HNSC. Asterisks indicate FDR-corrected significance: q < 0.05, q < 0.01, q < 0.001. Color scale is Spearman ρ. Abbreviations: RPPA, reverse-phase protein array; FDR, false-discovery rate.

#### 2.3.6 Protein-level mapping of pathway dependency

We mapped 54 RPPA proteins to their cognate pathways and correlated each protein with the predicted dependency of its own pathway across 212 TCGA-HNSC patients (proteins classified as drug targets, activity readouts, upstream regulators, or suppressors). Raw Spearman correlations identified several significant associations, the strongest being CASPASE7-cleaved (ρ = 0.48), BCL2 (0.32), and BIM (0.27) with APOPTOSIS dependency; CRAF (0.27) and CRAF pS338 (0.18) with MAPK; CYCLIN B1 (0.32) with CELL_CYCLE; HER2 pY1248 (0.26) with EGFR; and PTEN (0.20) with PI3K/MTOR (Fig. 6C, Table S10).

Because the nine predictions co-vary strongly, we recomputed each correlation as a partial correlation controlling for the other eight pathway dependencies. Under this adjustment, eight of nine pathways lost all protein-level signal. Only EGFR retained pathway-specific associations with HER2 pY1248 (ρ_raw = +0.26 → ρ_partial = +0.35, q < 0.001), HER3 (ρ_raw = +0.18 → ρ_partial = +0.25, q = 0.006), and EGFR pY1068 (ρ_raw = +0.02 → ρ_partial = +0.23, q = 0.009), and CYCLIN E1 retained a pathway-specific association with CELL_CYCLE dependency (ρ_raw = 0.17 → ρ_partial = +0.27, q = 0.002). No target protein for PI3K/MTOR, MAPK, APOPTOSIS, WNT, P53, JNK/P38, or IGF1R reached pathway-specific significance at q < 0.10 (Table S22). A consistent pattern emerged: direct drug-target proteins showed essentially no correlation with their own pathway’s dependency (PI3Kp110α: ρ = −0.11; AKT: −0.16; MTOR: −0.08; CDK1: +0.06; MEK1: −0.01; ERK2: −0.08; EGFR: −0.08; HER2: −0.15; GSK3α/β: −0.24; P53: −0.02), whereas the proteins that correlated most strongly were activity readouts and pathway suppressors (cleaved caspase-7, HER2 pY1248, PTEN, IRS1). The cross-pathway protein × dependency matrix confirmed this structure: most proteins correlated positively with their own pathway dependency and with the other eight, consistent with a shared underlying state, and only the EGFR proteins (HER2 pY1248, HER3, EGFR pY1068) showed a pathway-restricted pattern.

### 2.4 Prognostic value of the dependency signature

We tested whether the global dependency score was associated with progression-free interval (PFI) in TCGA-HNSC. For each patient, we computed a leave-one-out global signature (the mean of the eight remaining standardized pathway predictions) to avoid circularity. In HPV-stratified Cox regression, the global signature was not significantly prognostic in either stratum: HPV-positive (n = 39, 5 events, HR = 0.94, 95% CI 0.40–2.18, p = 0.88) or HPV-negative (n = 75, 28 events, HR = 0.92, 95% CI 0.63–1.35, p = 0.68). A PCA-weighted version of the global signature, in which the first principal component captured 55% of the variance across the nine pathway predictions, produced the same null result (HPV-positive HR = 0.94, 95% CI 0.39–2.25, p = 0.89; HPV-negative HR = 0.84, 95% CI 0.58–1.24, p = 0.39).

We next tested an HPV-stratified composite risk score, derived from univariate screening at p < 0.20 followed by multivariate Cox regression and applied as frozen coefficients to the independent GSE65858 cohort using overall survival as the endpoint. Univariate screening retained two pathways in each stratum: EGFR and MAPK in HPV-positive tumors; APOPTOSIS and CELL_CYCLE in HPV-negative tumors. In TCGA, the composite was significant in the HPV-positive stratum (HR = 3.45, 95% CI 1.05–11.3, p = 0.042; 5 events, underpowered) and in the HPV-negative stratum (HR = 1.54, 95% CI 1.06–2.25, p = 0.024). Neither signal was replicated in GSE65858 (HPV-positive HR = 1.17, 95% CI 0.71–1.93, p = 0.53; HPV-negative HR = 0.86, 95% CI 0.69–1.08, p = 0.20), and the HPV-negative stratum changed direction between cohorts. Kaplan–Meier analysis did not show significant separation between high- and low-risk groups in either GSE stratum (log-rank p > 0.05).

## 3. Discussion

In this study, we developed ElasticNet regression models to predict pathway-specific drug dependency using 50 pathway activity scores across 953 GDSC cell lines, based on negated, mean-per-drug, z-scored LN-IC50 values. The main goal was to distinguish “pathway activation” from “pathway dependency” in HNSCC patients using their genomic information. We trained nine models with CV R² ranging from 0.17 (IGF1R) to 0.48 (P53), with a median of 0.3 across pathways (Table 1, Figure 1A). We then applied them to HNSCC tumors in TCGA-HNSC and GSE65858, validating the predictions across genomic, pharmacological, and proteomic layers. Strikingly, IGF1R was the most structurally distinct pathway among the nine tested, with the lowest PC1 loading (0.167), the smallest leave-one-out global-signature correlation (ρ = 0.28, p = 6 × 10⁻¹²), and the only pathway with a significant negative co-dependency (IGF1R↔EGFR, ρ = −0.15, q = 3.2 × 10⁻⁴). It also consistently had the lowest CV R² (0.144 ± 0.016; rank 9/9 across 5 seeds). These results align with the performance of predictive models for complex traits and drug response, which generally have R² values of 0.01 to 0.3 (19, 20). The drug count showed no significant correlation with the IGF1R’s CV R². It was the best-calibrated model (slope = 0.955, r = 0.93) and the only one to benefit from pairwise interactions (ΔR² = +0.020 vs. negative ΔR² for all eight others), with a learning curve that continued to rise at full data.

Notably, IGF1R activation in HNSCCs occurs through a matrix-adhesion receptor complex consisting of syndecan-1 and αvβ3/αvβ5 integrin, present in >95% of HNSCCs but absent in benign epithelium, and is independent of the IGF1 ligand (21). This activation mechanism would not be simply inhibited by ligand-blocking antibodies, whose failure in HNSCC clinical trials is consistent with a dependency biology that does not align with the canonical ligand-driven framework on which both the drugs and our training target were based (22). IGF1R grouped with MAPK and EGFR into a minor secondary module in the two-factor EFA (75.3% cumulative variance; Factor 2 loadings: EGFR 0.88, MAPK 0.69, IGF1R −0.09) and was the opposing pole of the second principal component (IGF1R −0.50 vs. EGFR +0.60, MAPK +0.40).

IGF1R↔MAPK coupling was weak at the raw level (ρ = +0.10). This is consistent with the IGF1R→MAPK axis not being direct in the HNSCC context. Conversely, IGF-1-stimulated ERK activation requires EGFR kinase activity, and EGFR inhibition completely abrogates IGF-1-induced ERK phosphorylation while leaving Akt activation intact (23).

Our analyses show that, at the patient level, the dependency architecture is a single dominant axis rather than a set of modules. Pathways that load strongly on this axis and form the core dependency state are WNT, JNK_P38, PI3K_mTOR, CELL_CYCLE, and APOPTOSIS. However, this is a patient-level axis; given the variety of pathway dependencies at the cell level, the result is expected. Bulk data cannot resolve whether the co-dependency reflects a shared transcriptional state across cells, a compositional effect from the cell-type mixture within each tumor, or both. Consistent with the compositional interpretation, we found that the dominant axis correlates with CD8⁺ T-cell infiltration at ρ = 0.80, and CD8⁺ adjustment reduces PC1 variance from 62% to 48% while leaving a residual axis that is not stably estimable. The axis is therefore reproducible across cohorts and platforms, but a substantial fraction of its variance reflects immune infiltration rather than pathway-intrinsic co-dependency. However, coupled co-dependencies are more interesting from a therapeutic perspective; offering combination therapies for pathways on parallel signaling arms with positive correlation may be beneficial. Accordingly, targeting both PI3K/MTOR and MAPK (ρ=0.70 in HPV+ and ρ=0.67 in HPV-) may be synergistic, consistent with the evidence. Combination therapy with Tipifarnib (MAPK inhibitor) and Alpelisib (PI3K/mTOR inhibitor) showed synergistic antitumor effects in mouse models of HNSCC; however, their mechanisms differed between HPV+ and HPV– subgroups. In HPV+ models, Tipifarnib efficacy is CD8+ T-cell dependent, whereas in HPV-negative ones, the combination works primarily by overcoming intrinsic AKT-mediated resistance (24). Moreover, concurrently inhibiting vertically correlated pathways may help overcome resistance. For instance, the PI3K/MTOR↔CELL_CYCLE pair (ρ = 0.87 in HPV-positive, 0.69 in HPV-negative) represents a vertical pathway in which the downstream node (cell cycle) becomes a resistance mechanism when the upstream node (PI3K) is blocked. In PIK3CA-mutant HNSCC, inhibiting PI3K leads to sustained Rb phosphorylation via AKT-independent mTORC1 activation. Adding a CDK4/6 inhibitor (Abemaciclib) or an mTORC1 inhibitor (Everolimus) can further restore sensitivity in resistant preclinical HNSCC models (25).

The RPPA-measured abundance of the direct drug-target protein (PI3Kp110α, AKT, MTOR, CDK1, MEK1, ERK2, EGFR, HER2, GSK3α/β, P53) across all pathways showed no correlation with the predicted dependency of its own pathway (all |ρ| < 0.20; most between −0.15 and +0.06). This further underscores the distinction between target-protein abundance and pathway dependency and may explain the failure of targeted-therapy response prediction based on target expression level (9, 26–29). In contrast, we tracked dependency using readouts of pathway-suppressor proteins, including cleaved caspase-7 for APOPTOSIS (ρ = 0.48), HER2 pY1248 for EGFR (ρ = 0.26), PTEN for PI3K/MTOR (ρ = 0.20), and IRS1 for IGF1R (ρ = −0.22). However, in vivo and in vitro studies are needed to further assess this claim. After adjusting for covariates, seven pathways showed significantly higher dependency in HPV+ patients across both the TCGA and GSE65858 datasets: P53, PI3K/MTOR, WNT, IGF1R, JNK/P38, APOPTOSIS, and CELL_CYCLE. While the model predicted higher P53 dependency in HPV+ tumors, the HPV E6 oncoprotein has been shown to cause p53 degradation (14). Notably, the model predicts dependency on the hallmark P53 transcriptional program rather than on p53 protein abundance. The hallmark P53 set includes 190 genes, including TP53. Our results show that none of these genes abolished the HPV→P53 dependency association after FDR correction, indicating that the effect is distributed across the program rather than concentrated in a single node. Moreover, genes upregulated in HPV-positive tumors were predominantly p53 transcriptional targets (POLH, TP53, ISCU, XPC, CYFIP2, TSPYL2), whereas genes downregulated in HPV-positive tumors were predominantly p53-repressed (SLC7A11, SLC3A2, UPP1, SDC1, TGFA, HBEGF, KLK8, KRT17), highlighting that p53 is activated at the transcriptional level. By computing a p53 target score (mean expression of 13 canonical p53 targets) and a p53 repressor score (mean expression of 12 p53-repressed genes), we derived a net p53 activity score that was 2.5 log units higher in HPV-positive than in HPV-negative tumors (r = 0.75, q < 0.001). Critically, this signature was stronger in the TP53-WT stratum than in the full cohort, indicating that the TP53-mutant minority does not inflate the association.

Intriguingly, the p53-active transcriptional signature in TP53-WT tumors aligns with evidence that HPV does not fully inactivate p53 in HNSCC; rather, residual, tumor-suppressive p53 activity is retained. siRNA- and CRISPRi-mediated p53 depletion in these cells increased proliferation, migration, and invasion, and transcriptomic analysis confirmed ongoing p53-dependent gene regulation despite the presence of HPV (30). According to our results, TP53 mRNA was also highest in HPV-positive/TP53-WT tumors (median 11.41 vs. 10.36 in HPV-negative/TP53-WT, p < 0.0001) and correlated with the p53 target score (Spearman ρ = 0.417, 95% CI 0.05–0.68, ρ² = 0.17, p = 0.027), consistent with the mechanism originally described in normal cervical keratinocytes, in which E6-mediated degradation of p53 protein is compensated by a 1.5- to 2-fold upregulation of TP53 mRNA, thereby maintaining p53 protein levels (31). The ∼2-fold increase in TP53 mRNA in HPV+/TP53-WT tumors is quantitatively similar to the 1.5- to 2-fold upregulation reported by Kawamata et al. in E6-expressing cervical keratinocytes; however, the difference in context warrants attention.

PI3K/MTOR was the most confounded pathway in our model, and its prediction correlated with CD8⁺ at ρ = 0.85 in TCGA and ρ = 0.78 in GSE65858. Mediation analysis across the Hallmark PI3K/AKT/MTOR gene set failed to identify a pathway-specific signal (empirical p = 0.29), with the top attenuators dominated by immune signaling genes, and CD8⁺ adjustment reduced the HPV association by 72% (β = +0.29 → +0.08). The residual β = +0.08 (q = 0.015) is therefore unlikely to reflect a PI3K-specific biological effect. Evidence from the literature points in the opposite direction: INPP5D is upregulated in HPV+/TP53-WT tumors and suppresses PI3K-AKT signaling (1, 30), suggesting that true biological PI3K activity is lower, not higher, in HPV-positive tumors. The apparent higher PI3K/MTOR dependency in our model is therefore best explained by the confounding described in Section 2.2.2 rather than by a PI3K-specific biological finding.

The HPV-associated increase in predicted PI3K/MTOR dependency captured in our model was not attributable to the Hallmark PI3K/AKT/MTOR gene set: mediation analysis failed to identify pathway-specific signal (empirical p = 0.29), and the top attenuators were predominantly immune signaling genes. CD8 adjustment substantially attenuated the association (β = +0.29 to +0.08, a 72% reduction), suggesting that a large fraction of the HPV→PI3K/MTOR dependency association in patient tumors is immune-mediated. A small residual component (β = +0.08, q = 0.015) remained after CD8⁺ adjustment, which may reflect a smaller PI3K-specific contribution or residual confounding by other immune features not captured by the CD8⁺ marker.

Despite different definitions of dependency and activation, HPV+ tumors show enhanced proliferation. E7 promotes cell proliferation by binding to pRB, thereby enhancing the transcription factor E2F and increasing cell-cycle progression (32). Moreover, the proliferation index (Ki-67) is elevated in HPV+ patients (33). These are consistent with the positive dependency our results suggest for HPV+ tumors on the CELL_CYCLE pathway.

P53 degradation in HPV+ tumors leads to evasion of apoptosis in the cells (34, 35); however, in a large number of primary head and neck cancers and in the absence of functional p53, the apoptotic pathway is still active through other mechanisms such as Noxa induction via ATF3/ATF4 transcription factors (36), p73-mediated activation of pro-apoptotic targets (37, 38), BH3-mimetic-induced mitochondrial apoptosis (38), and death receptor signaling through caspase-8 that is unaffected by p53 restoration (39). Combination targeting of each of them has led to enhanced apoptosis regardless of HPV status in head and neck cancers (36–39), demonstrating cancer’s dependency despite probable less activation of this pathway in HPV+ HNSCC tumors.

After CD8+ adjustment, MAPK and EGFR shifted to negative coefficients (β = −0.39 and −0.43, q < 0.001), while JNK/P38 remained positive (β = +0.10, q = 0.015). All three pathways showed moderate CD8+ confounding (ρ = 0.38–0.70 across cohorts), and their CD8+-adjusted residuals should not be interpreted as direct HPV effects. The directions, however, remain consistent with known biology. EGFR activation suppresses type 1 cytokine-induced production of T-cell-attracting chemokines (CCL2, CCL5, CXCL9, CXCL10) via the MEK and JNK pathways (40); moreover, HPV+ tumors have lower expression of EGFR, pEGFRY1068, and pEGFRY1173 (41). HALLMARK_MAPK_SIGNALING primarily represents the canonical RAS-RAF-MEK-ERK cascade downstream of growth factor receptors, whereas HALLMARK_JNK_P38_PATHWAY captures the stress-activated JNK and p38 branches, which are activated by inflammatory cytokines and viral oncoproteins, in addition to EGFR. Evidence shows that HPV E6 activates JNK through its PDZ-binding motif, inducing c-Jun/AP-1 transcriptional activity and establishing a feed-forward loop that sustains viral oncogene expression (42). Ma *et al.* (40) also demonstrate that JNK inhibition had a stronger effect on HPV+ patients. Whether these mechanisms account for the adjusted residuals cannot be determined from cross-sectional data and would require experimental validation.

WNT dependency in HPV+ patients’ prediction correlated with CD8+ (ρ = 0.79 in TCGA and ρ = 0.72 in GSE); however, this association disappeared after adjustment (β = +0.17 unadjusted → −0.01 adjusted, q = 0.79). This suggests that immune infiltration drove the prediction, consistent with the inverse relationship between WNT activity and T-cell infiltration in HNSCC. The result exemplifies the confounding discussed in Section 2.2.2, not a WNT-specific biological finding.

This study has undeniable limitations: first, alcohol, despite its important role in the emergence of HNSCCs, was excluded from the primary model due to large missingness. Moreover, our immune adjustment used single immune markers and did not capture the full immune compartment, and we also did not include treatment. Second, the GDSC cell lines lacked immune cells. Third, we interpreted residual p53 based on transcriptional evidence rather than protein-level measurement. Some sample sizes, such as the HPV+ TP53-mutant group, were also too small; thus, the analyses were underpowered. The RPPA-based EGFR comparison by HPV was also not possible (n = 1 HPV+ tumor with RPPA data). Lastly, this study was a computational assessment with no experimental validation.

## 4. Materials and Methods

### 4.1 Data sources

All analyses were conducted in Python 3.11 using the following libraries: pandas, numpy, scikit-learn, scipy, statsmodels, gseapy, lifelines, networkx, matplotlib, seaborn, and joblib.

#### 4.1.1 Drug response data (GDSC)

Fitted dose–response curves from the Genomics of Drug Sensitivity in Cancer (GDSC) project were downloaded for both screening panels: GDSC1_fitted_dose_response_15Oct19.xlsx and GDSC2_fitted_dose_response_15Oct19.xlsx. The two tables were concatenated into a single dataframe (gdsc_full) and restricted to the columns SANGER_MODEL_ID, DRUG_ID, and LN_IC50 (natural-log IC₅₀). Duplicate drug–cell line pairs present in both panels were averaged before downstream processing. For the drug-holdout analysis (see below), CELL_LINE_NAME was also retained.

#### 4.1.2 Compound annotation and pathway mapping

Drug metadata were obtained from screened_compounds_rel_8.1.csv, which provides DRUG_ID, DRUG_NAME, and TARGET_PATHWAY. The nine target pathways used throughout were defined by the following mapping of the source TARGET_PATHWAY categories: PI3K/MTOR signaling: PI3K_MTOR; ERK MAPK signaling: MAPK; EGFR signaling: EGFR; WNT signaling: WNT; Apoptosis regulation: APOPTOSIS: Apoptosis; Cell cycle: CELL_CYCLE; Mitosis: CELL_CYCLE; IGF1R signaling: IGF1R; JNK and p38 signaling: JNK_P38; p53 pathway: P53.

Drugs whose TARGET_PATHWAY did not appear on this list were excluded, yielding a final set of nine dependency targets. A dictionary compound_to_pathway mapping DRUG_ID → pathway was constructed and applied to gdsc_full to produce gdsc_full_pathway, retaining only drug–cell-line pairs with an assigned pathway.

#### 4.1.3 Gene expression (GDSC cell lines)

Baseline transcriptomic profiles of GDSC cell lines were obtained from RNA_Seq_All_TPM.csv (columns: gene_symbol, model_id, rsem_tpm). The table was pivoted into a gene × cell-line matrix (expr_full) using pivot_table (index=’gene_symbol’, columns=’model_id’, values=’rsem_tpm’). Duplicate gene symbols were resolved by keeping the first occurrence. The resulting matrix contained 41,143 genes × 1,898 cell lines.

#### 4.1.4 Gene sets

The 50 MSigDB Hallmark gene sets were extracted from Human MSigDB Hs Symbols v2026.1.gmt by filtering lines whose first tab-delimited field began with HALLMARK_, retaining the gene list from field 3 onward.

#### 4.1.5 TCGA-HNSC cohort

The TCGA Head and Neck Squamous Cell Carcinoma (HNSC) cohort was used as the discovery cohort. The following files were used: phenotype.txt (clinical annotations), tcga_survival_ready_for_pfi.csv (progression-free interval; columns PFI, PFI.time), tcga_expression_ready_for_gsva.csv (expression matrix for ssGSEA), tcga_50pathway_scores.csv (pre-computed Hallmark ssGSEA scores), Gistic2_CopyNumber_Gistic2_all_thresholded.by_genes (thresholded gene-level copy-number calls), RPPA_RBN.txt (reverse-phase protein array), HNSC_mc3_gene_level.txt (binary gene-level mutation matrix), and HNSC_mc3.txt (mutation annotation file, MAF-style).

#### 4.1.6 External validation cohort (GSE65858)

GSE65858 (N = 270 HNSCC) was used as an independent external validation cohort. Expression was obtained from GSE65858_expression_genes.csv; HPV status was parsed from GSE65858_family.soft.gz using the Sample_characteristics_ch1 field (hpv_dna), classifying HPV16, HPV18, HPV16/18, POSITIVE, and POS as HPV-positive and NEGATIVE/NEG as HPV-negative. Survival data were obtained from gse65858_survival.csv (OS, OS.time).

#### 4.1.7 Effective sample sizes across analyses

Analyses had slightly different effective sample sizes depending on data availability. The HPV-status comparison required only HPV annotation (n = 114; 39 HPV+, 75 HPV−). Covariate-adjusted OLS models required complete covariate data (n = 111; 39 HPV+, 74 HPV−). Co-occurrence network analyses and subgroup comparisons used the same respective subsets. Survival analyses required PFI and used the full set with available predictions and clinical annotation (n = 114 for the global signature analysis; n = 114 for the composite risk-score derivation, 39 HPV+ and 75 HPV−). These differences are noted where relevant in the Results.

### 4.2 Preprocessing and target definition

#### 4.2.1 IC50 missingness analysis

Because GDSC does not test every drug in every cell line, we first tested whether IC50 missingness was random or informative. For each drug, we compared the pathway dependency score between cell lines in which the drug was tested versus those in which it was not.

For each “cell line, pathway” pair, a leave-one-drug-out dependency score was computed as the mean per-drug z-scored LN-IC50 across all other drugs in that pathway, excluding the drug under consideration. Drugs with fewer than 20 tested cell lines were discarded. For each retained drug, we defined:

- **Group 1**: cell lines with observed IC50 for that drug;
- **Group 2**: cell lines with missing IC50.

A two-sided Mann–Whitney U test was applied, requiring ≥5 samples per group. p-values were adjusted using the Benjamini–Hochberg false-discovery-rate (BH-FDR) procedure; associations with q < 0.05 were considered significant.

#### 4.2.2 Dependency scores

After confirming that missingness was largely non-informative or affected only a small number of pairs, missing values in the raw dependency matrix were imputed with the median per pathway. Scores were first averaged across any duplicate drug–cell line pairs introduced by concatenating GDSC1 and GDSC2, then z-scored within each drug across all pathway-assigned cell lines:

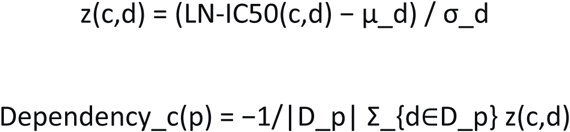

where μ_d and σ_d are the mean and standard deviation of LN-IC50 for drug d across all cell lines in gdsc_full_pathway. The final matrix contained 978 cell lines × 9 pathways.

#### 4.2.3 Feature construction (GSVA)

Before pivoting, the RNA-Seq table was restricted to the union of genes present in the 50 MSigDB Hallmark gene sets. The resulting Hallmark-restricted gene × cell-line matrix was used for ssGSEA. Duplicate gene symbols were resolved by keeping the first occurrence. ssGSEA was run with method=’ssgsea’, min_size=5, max_size=1000, and parallel=False. The resulting feature matrix contained one enrichment score per Hallmark set per cell line.

### 4.3 Model training

#### 4.3.1 Elastic Net modeling training

Predictive models for the nine dependency targets were trained with Elastic Net regression using scikit-learn’s ElasticNetCV. For each target pathway, we performed 5-fold outer cross-validation (KFold(n_splits=5, shuffle=True, random_state=42)). Within each fold:

- Features were standardized (StandardScaler) on the training fold only, and the same scaler was applied to the validation fold.
- Hyperparameters were tuned with an inner 5-fold cross-validation using the grid l1_ratio ∈ {0.1, 0.3, 0.5, 0.7, 0.9, 0.95, 0.99, 1.0} × α ∈ logspace(−4, 1, 50), max_iter=10000, n_jobs=−1.
- The tuned model was refit on the training fold and used to predict the validation fold.

Model performance was summarized as the mean R² across the five outer folds.

To prevent information leakage, per-drug means and standard deviations for z-scoring LN-IC50 were calculated using only training-fold cell lines, then applied to validation-fold lines. Pathway dependency targets were reconstructed within each fold using these statistics, ensuring validation targets never influenced feature or drug-response scaling.

#### 4.3.2 Interaction screening

For each of the nine dependency targets, an ElasticNetCV model was fit on the full 953-cell-line dataset using all 50 Hallmark GSVA features. In the current implementation, all pairwise multiplicative products of the 50 features were generated, yielding 1,225 interaction terms.

Under the same 5-fold outer CV scheme, two models were compared per fold: Base model: ElasticNetCV trained on the 50 original features.

Interaction model: ElasticNetCV trained on the 50 original features plus all 1,225 pairwise products.

Features were standardized on the training fold before model fitting. R² was computed on the validation fold and averaged across folds. Improvement was defined as ΔR² = mean R²(interaction) − mean R²(base). A target was considered to benefit from interactions if ΔR² > 0.01 (43).

#### 4.3.3 Final models

For pathways meeting ΔR² > 0.01, final interaction models were retrained using the same 5-fold CV scheme, standardization, inner-CV hyperparameter search, and interaction construction described above. For remaining pathways, the additive 50-feature model was retained.

#### 4.3.4 Bootstrap stability analysis

Parameter uncertainty was estimated by non-parametric bootstrapping. For eight pathways, 1,000 bootstrap resamples were used; for IGF1R, 500 resamples were used due to computational cost. For each pathway, the model α and l1_ratio were fixed to the values selected on the full data, and an ElasticNet model was refit on each bootstrap resample of the 953 cell lines (sampling with replacement). The intercept 95% CI, mean coefficient, coefficient 95% CI, and selection probability were recorded. Selection probability was defined as the fraction of bootstraps in which the coefficient was non-zero. Features with selection probability > 0.5 were considered stable.

#### 4.3.5 Calibration analysis

For each pathway, out-of-fold predictions were collected using the same leakage-free 5-fold CV scheme. Predictions were binned into deciles, and mean predicted vs mean observed values were computed per bin. A calibration slope was fit as observed ∼ predicted across bins. A slope < 1 indicates over-confidence; > 1 indicates under-confidence; ∼1 indicates good calibration. Pearson correlation between binned predicted and observed values was also reported.

#### 4.3.6 Drug-count vs CV R^2^

To test whether pathway model performance was explained by the number of drugs used to construct the dependency target, the number of drugs per pathway was computed from screened_compounds_rel_8.1.csv. Spearman and Pearson correlations between drug count and CV R² were computed. A linear regression of CV R² on drug count was fit, and the residual R² for IGF1R was reported.

#### 4.3.7 IGF1R stability analysis

Three tests were performed:

- Repeated CV: 5-fold CV was repeated across five random seeds. IGF1R’s rank among the nine pathways was recorded.
- Interaction benefit: ΔR² from interaction models was compared across all nine pathways.
- Learning curve: Models were trained on 25%, 50%, 75%, and 100% of training cell lines, and CV R² was computed for each fraction. The R² gain from 75% to 100% was used to assess whether IGF1R was data-limited.

### 4.4 Internal validation

#### 4.4.1 Leave-One-Pathway-Out (LOPO) validation

For each held-out pathway P:

1. Training drugs were those annotated to any pathway other than P.
2. Held-out drugs were those annotated to P.
3. For each outer fold (5-fold CV on cell lines), a training dependency score was computed as the mean per-drug z-scored LN-IC50 across training drugs, negated.
4. A held-out dependency score was computed as the mean per-drug z-scored LN-IC50 across held-out drugs, with drug means/SDs estimated from training cells only, negated.
5. An ElasticNetCV model was trained on the 50 Hallmark features of the training cell lines and used to predict held-out dependency on the test cell lines. Spearman’s ρ was the primary metric; R^2^ was secondary.

#### 4.4.2 Drug–pathway correlation analysis

For each drug with ≥10 measured cell lines, the mean LN-IC50 per cell line was correlated with each pathway’s dependency score using Spearman ρ. For the drug’s own target pathway, dependency was computed leave-one-drug-out. All other pathway columns were unchanged. 95% CIs were computed via Fisher Z. P-values were BH-FDR-corrected across all drug–pathway pairs. Significant negative correlations were defined as ρ < −0.3 and q < 0.05.

#### 4.4.3 Drug-target validation (top-25% dependency)

For each drug, target-pathway dependency scores and 75th-percentile cutoffs were calculated using leave-one-drug-out. Cell lines were classified as High (≥75th percentile) or Low (<75th). When the drug’s target pathway matched the pathway studied and each group contained ≥5 cell lines, LN-IC50 values were compared using a one-sided Mann–Whitney U test (alternative=’less’). Effect size r, the Hodges–Lehmann estimate, and bootstrap 95% CIs were calculated; P-values were BH-FDR-corrected.

### 4.5 TCGA-HNSC inference and clinical associations

#### 4.5.1 TCGA-HNSC inference

Final GDSC models were refit on the full 953-cell dataset after fitting StandardScaler on the same matrix. Both models and the scaler were persisted. Pre-computed Hallmark ssGSEA scores for TCGA-HNSC (tcga_50pathway_scores.csv) were re-ordered to match the training feature order, standardized using the frozen scaler, and passed through the nine models to produce predicted dependency scores. Models were trained on the per-drug z-scored dependency target, so the resulting predictions are on that scale (mean near zero across the training set) Clinical annotations were extracted from phenotype.txt, restricted to patients with available predictions.

The following covariates were defined: age (age_at_initial_pathologic_diagnosis), gender (1 for MALE, 0 for FEMALE), stage (ordinal 1–4 from clinical_stage, mapping Stage IVA/B/C and IV to 4), smoking (0 (never), 1 (former), 2 (current/reformed), from tobacco_smoking_history), alcohol (0, 1, 2 from frequency_of_alcohol_consumption), HPV (1 for positive/pos (from hpv_status_by_p16_testing), 0 for negative/neg), and treatment_Score (sum of surgery, radiation, and chemotherapy indicators).

#### 4.5.2 HPV association

For each pathway, an OLS regression was fit: “Dependency ∼ HPV + Age + Gender + Stage + Smoking + Treatment_Score”

Continuous covariates were standardized. Alcohol was not included in the final multivariate model; it was analyzed separately in phenotype associations. Complete-case N = 111 (38 HPV+, 73 HPV−). A univariate Mann–Whitney U test (HPV+ greater than HPV−) was also reported on the same complete-covariate subset. Both p-values were BH-FDR-corrected across the nine pathways.

#### 4.5.3 HPV association adjusted for CD8⁺ infiltration

To isolate the direct effect of HPV on predicted dependency from the confounding effect of immune infiltration, we fit, per pathway, the OLS model:

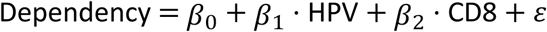

The CD8⁺ score was computed as the mean expression of five canonical cytotoxic T-cell markers (CD8A, CD8B, GZMA, GZMB, PRF1) when all were present in tcga_expression_ready_for_gsva.csv; if fewer than one of these genes was available, the HALLMARK_ALLOGRAFT_REJECTION ssGSEA score was used as a proxy.

The HPV coefficient β₁, its 95% CI, and its p-value were extracted and BH-FDR-corrected them across the nine pathways.

A sensitivity analysis was performed to test whether the CD8-adjusted HPV effects on EGFR and MAPK were confounded by smoking, which is enriched in HPV-negative HNSCC and known to activate RTK signaling. Smoking status (0 = never, 1 = former, 2 = current/reformed) was added to the CD8-adjusted model, with the sample held constant (N = 112).

#### 4.5.4 P53/PI3K per-gene mediation and mutation stratification

For each Hallmark P53 or PI3K/AKT/MTOR gene, a baseline OLS model was fit:

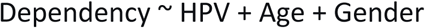

Dependency ∼ HPV + Age + Gender Then a mediation model was fit:

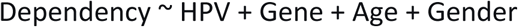

Attenuation was defined as (β_base − β_adj) / β_base. P-values for the gene term and adjusted HPV term were BH-FDR-corrected across genes. Candidate mediator genes met all three criteria: r_gene_HPV > 0, q_gene < 0.05, and q_HPV_adj > 0.05.

A circularity check was performed by repeating the mediation analysis on 500 random gene sets sampled from non-pathway genes matched for mean expression. Median and maximum attenuation were compared between real and random gene sets.

For TP53 stratification, patients were classified as TP53-WT or TP53-mut. The HPV effect on P53 dependency was estimated within All, TP53-WT, and TP53-mut strata using:

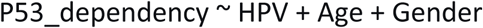

p53 transcriptional scores were computed as the mean expression of p53 target genes minus the mean expression of p53-repressed genes. HPV+ vs HPV− differences were tested by Mann– Whitney U within strata, with BH-FDR correction.

For PI3K, analogous stratification was performed by PIK3CA mutation, PTEN mutation, PIK3CA amplification, and PTEN deletion.

#### 4.5.5 Treatment effect

Patients were grouped into High (Treatment_Score ≥ 2), Low (= 1), and None (= 0). Mann– Whitney U tests (two-sided) compared High vs Low and High vs None per pathway. All p-values across both comparisons and all pathways were pooled and BH-FDR-corrected.

#### 4.5.6 Phenotype associations

For each of {Age, Stage, Smoking, Alcohol} and each pathway, a Spearman correlation was computed with ≥20 paired observations and BH-FDR-corrected across all tests.

#### 4.5.7 Stage association

A Kruskal–Wallis test compared dependency across ordinal stage groups per pathway (≥10 observations, ≥2 unique stages) with BH-FDR correction. Box plots for the top three pathways by lowest p-value were generated.

### 4.6 TCGA genomic and proteomic validation

#### 4.6.1 Copy-number validation

Thresholded gene-level copy-number calls (Gistic2_CopyNumber_Gistic2_all_thresholded.by_genes) were aligned with predictions after detecting the sample axis by matching a TCGA barcode regex. Four a priori hypotheses were evaluated: EGFR amplification → EGFR dependency, PTEN deletion → PI3K_MTOR dependency, CDKN2A deletion → CELL_CYCLE dependency, and CCND1 amplification → CELL_CYCLE dependency.

For each hypothesis, patients were split into Altered (CNV ≥ +1 for amplifications, ≤ −1 for deletions) and Normal (CNV = 0) groups. Predicted dependency was compared using a two-sided Mann–Whitney U test (asymptotic). Additional statistics included: 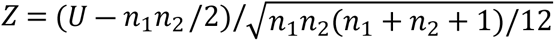; effect size 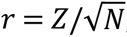; the probability of superiority PSet = *U*/(*n*_1_*n*_2_); and the Hodges–Lehmann estimate (median of all pairwise cross-group differences) with a bootstrap 95% CI (1,000 resamples). p-values were BH-FDR-corrected across the four hypotheses.

#### 4.6.2 Gene-level mutation validation

A binary gene-level mutation matrix (HNSC_mc3_gene_level.txt) was transposed to samples × genes and merged with predictions. For each predefined gene–pathway pair (TP53 → P53, PIK3CA → PI3K_MTOR, KRAS → MAPK, BRAF → MAPK, EGFR → EGFR, PTEN → PI3K_MTOR), predicted dependency was compared between mutant (≥3 samples) and wild-type patients using a two-sided Mann–Whitney U test. Effect sizes (Z, r), the Hodges–Lehmann estimate, and its bootstrap 95% CI were computed as for the CNV analysis. p-values were BH-FDR-corrected.

#### 4.6.3 Variant-class mutation validation

Mutation calls from HNSC_mc3.txt were used to classify variants by functional impact. Columns were renamed to Tumor_Sample_Barcode, Hugo_Symbol, Variant_Classification, and HGVSp_Short; samples were restricted to TCGA barcodes. Per-gene classification rules:

- TP53: LoF (Nonsense, Frameshift, Splice_Site, Translation_Start_Site); Hotspot (Missense containing R175H, G245S, R248Q/W, R249S, R273H/C, or R282W); Missense (other); Indel (In_Frame_Del/Ins); Other.
- BRAF: V600E (Missense containing V600); Other (Missense or In_Frame); LoF (Nonsense, Frameshift, Splice_Site).
- PIK3CA: Hotspot (Missense containing E542K, E545K/A/G, Q546K, H1047R/L/Y); Missense; LoF; Indel; Other.
- PTEN: LoF; Missense; Indel; Other.

When a patient carried multiple mutations in the same gene, the most severe class was retained using an ordinal severity ranking (LoF > Hotspot > Missense > Indel > Other for TP53; V600E > Other > LoF for BRAF; etc.). Wild-type patients were those without any variant in that gene.

For each (gene, mutation class, pathway) triplet with ≥5 mutant samples, predicted dependency was compared to wild-type patients using the same statistics as the gene-level analysis. p-values were BH-FDR-corrected across all tests.

#### 4.6.4 RPPA single-protein validation

RPPA data (RPPA_RBN.txt) were transposed to samples × proteins. A curated protein-to-pathway dictionary was defined.

For each available protein and its target pathway, Spearman ρ between protein (or phosphoprotein) abundance and predicted dependency was computed with ≥10 paired samples. 95% CIs were derived via Fisher’s Z-transformation. A secondary Mann–Whitney U test compared predicted dependency between patients with protein above vs. below the median. p-values were BH-FDR-corrected across all protein–pathway pairs.

#### 4.6.5 RPPA-based co-activity analysis

For each pathway, a directional protein-activity score was computed as the mean z-scored activator proteins minus the mean z-scored suppressor proteins. This prevents functionally opposing proteins from canceling. Pairwise Spearman correlations between pathway activities were computed with Fisher Z 95% CIs and BH-FDR correction (|ρ| > 0.2, q < 0.10). The RPPA correlation matrix was compared with the prediction-based co-dependency matrix.

#### 4.6.6 Hallmark signature correlation (TCGA)

Single-sample GSVA was performed on tcga_expression_ready_for_gsva.csv using eight selected Hallmark gene sets (HALLMARK_PI3K_AKT_MTOR_SIGNALING, HALLMARK_P53_PATHWAY, HALLMARK_EPITHELIAL_MESENCHYMAL_TRANSITION, HALLMARK_G2M_CHECKPOINT, HALLMARK_APOPTOSIS, HALLMARK_MYC_TARGETS_V1, HALLMARK_TNFA_SIGNALING_VIA_NFKB, and HALLMARK_HYPOXIA). The implementation used a rank-based ssGSEA with min_size=5 and max_size=500. Spearman correlations were computed between every signature and every predicted dependency. P-values were BH-FDR-corrected.

#### 4.6.7 IGF1R downstream validation

IGF1R dependency was correlated with canonical RPPA readouts (IRS1, p-AKT T308/S473, p-MEK, p-ERK, p-MTOR, p-P70S6K, p-4EBP1) using Spearman correlation with BH-FDR. Reverse checks correlated p-AKT and p-ERK with all nine dependency pathways. In GSE65858 and TCGA, IGF1R dependency was correlated with MAPK, PI3K/AKT/MTOR, and IGF1R-axis gene-expression signatures.

### 4.7 Pathway co-dependency architecture (TCGA)

#### 4.7.1 Pathway co-occurrence analysis

Pairwise co-occurrence of predicted pathway dependencies was assessed on TCGA-HNSC predictions.

For every unordered pair of pathways, Spearman ρ was computed across patients with ≥3 unique values per pathway. The 95% CI was derived via Fisher’s Z-transformation with standard error 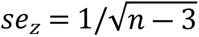. p-values were BH-FDR-corrected. A pair was designated significant if q < 0.05 and |ρ| > 0.3.

The procedure was repeated in four subgroups: HPV+, HPV−, early stage (I–II), and late stage (III–IV).

#### 4.7.2 Global dependency signature

For each patient, a leave-one-out (LOO) global dependency score was the mean predicted dependency across the other eight pathways. Calculating it separately for each pathway avoided self-correlation. Spearman correlation between each pathway and its LOO score measured alignment with the global axis.

Dimensionality was assessed by PCA of the z- scored nine- pathway matrix (TCGA-HNSC, n = 565). PC 1 variance was compared with a null from 1, 000 permutations that independently shuffled each pathway. Parallel analysis used the same scheme, retaining components with eigenvalues above the permuted 95th percentile. Bootstrap 95% CIs for PC 1 loadings came from 1, 000 patient resamples; PC 1 signs were aligned to the reference before calculating percentiles. Internal consistency was measured by Cronbach’s α. Factor structure was tested using 1-, 2-factor, and bifactor CFA models (ML estimator). Since absolute fit indices are unreliable when inter-item correlations exceed ∼0. 85, we assessed explained common variance (ECV), omega hierarchical (ωH), omega subscale (ωS), and omega total (ωT). A free 2-factor EFA with varimax rotation provided a comparison. Split-half stability was assessed across 100 random splits by fitting EFA to each half and calculating Tucker congruence between loading matrices. EFA loading stability was also assessed across 1, 000 bootstrap resamples, recording each pathway’s dominant-factor assignment frequency.

Modularity was assessed independently using exploratory graph analysis (EGA) of the partial-correlation network and community detection on raw correlations. Partial correlations came from the inverse correlation matrix; edges with |partial ρ| > 0. 05 were retained, and Louvain optimization detected communities. Raw-correlation communities were tested across thresholds (|ρ| ≥ 0. 2- 0.2- 0.7), with 500 bootstraps at |ρ| ≥ 0. 0.4. Co-membership stability was the fraction of bootstraps in which each of the 36 pathway pairs clustered together. The 2-factor EFA was also repeated on 100 random halves to assess loading stability.

To measure how much pairwise co-dependency the global axis explained, each pathway was regressed on PC 1, and Spearman correlations were recalculated on residuals. We report the median absolute reduction across 36 pairs and the number retaining | ρ _ resid | > 0. 0.3. For collinearity, each pathway was regressed by OLS on the other eight, and VIFs were calculated. For covariate adjustment, each pathway was regressed on HPV status, stage, age, and sex; PCA was repeated on residuals (n = 113 complete cases). Replication used GSE 65858 (n = 270): PCA was performed on z-scored pathway predictions, and PC 1 loadings were compared with TCGA using Tucker congruence.

#### 4.7.3 Protein-level dependency mapping (RPPA)

We examined whether the co-dependency observed in predicted scores reflected protein-level co-regulation. To construct RPPA pathway activity scores, we assigned each of the 54 RPPA proteins to one of the nine pathways and classified them by role (direct drug target (19), activity readout (21), upstream regulator (10), or pathway suppressor (4). Role assignment was fixed a priori and not adjusted based on results (Table S). For each pathway, the score was computed as the mean of z-scored activator proteins minus the mean of z-scored suppressor proteins, so that functionally opposing proteins (e.g., AKT and PTEN, BCL2 and BAX) contributed in opposite directions rather than canceling.

For the next analysis, samples were restricted to the 212 TCGA-HNSC patients with both RPPA measurements and predicted dependency scores. For each protein–pathway pair, we computed a Spearman correlation between the protein’s RPPA value and the predicted dependency of the matched pathway (Analysis A). To isolate pathway-specific signal from the shared dependency state, we recomputed each correlation as a partial correlation controlling for the predicted dependencies of the other eight pathways using linear regression residuals (Analysis B). All correlations were FDR-corrected across proteins by Benjamini–Hochberg.

Candidate dependency nodes were defined as proteins classified as direct drug targets with a partial correlation > +0.15 and q < 0.10 in Analysis B. Active-pathway readouts and lost suppressors were defined as readouts or suppressors with q < 0.05 and |ρ_raw| > 0.20 in Analysis A, for contextual interpretation (Analysis C).

A cross-pathway matrix (Analysis D) was constructed by correlating each of the 54 proteins against each of the nine pathway dependencies, giving a 54 × 9 matrix of Spearman ρ values. Proteins were grouped by role and displayed as a heatmap (RdBu_r colormap, ρ ∈ [−0.5, +0.5]) to visualize pathway-specific versus shared protein–dependency associations.

#### 4.7.4 Single-cell co-dependency analysis (GSE103322)

GSE103322 was processed by separating the first five metadata rows from the expression matrix. Cells were classified as malignant or non-malignant using the metadata flags. Malignant cells were retained, and the expression matrix was converted to numeric. Hallmark ssGSEA was run on malignant cells. Frozen GDSC ElasticNet models were applied to the Hallmark scores to produce single-cell pathway dependency predictions.

Pairwise Spearman correlations between pathway predictions were computed across malignant cells, with BH-FDR correction; significant pairs were defined as |ρ| > 0.3 and q < 0.05. Pseudobulk predictions were computed by averaging predictions per patient. Pairwise pseudobulk correlations were computed. Agreement between single-cell, pseudobulk, and bulk TCGA correlation matrices was assessed by Pearson correlation.

PC1 was computed on z-scored single-cell and pseudobulk predictions. PC1 loadings were compared to bulk TCGA PC1 loadings using Tucker congruence. Leave-one-out global dependency scores were computed as the mean of the other eight pathways. Parallel analysis used 200 permutations for single-cell and 500 for pseudobulk. Bootstrap PC1 95% CIs used 500 resamples. Residual correlations after removing PC1 were computed. Cronbach’s α, CFA (1-factor, 2-factor, bifactor), and EGA on partial-correlation networks were performed.

Null tests included:

- Random-weight null: 1,000 random weight matrices.
- Permutation null: each dependency target was shuffled, models were refit, and PC1 variance was recomputed (200 permutations).
- Cross-platform concordance from model weights was compared to 500 random-weight matrices.
- Dependency concordance after removing 5 Hallmark PCs was also computed.

### 4.8 Survival analyses (TCGA)

For each HPV stratum, pathway predictions were z-scored within the stratum. For each pathway, a univariate Cox proportional-hazards model was fit against progression-free interval (PFI). Pathways with p < 0.20 were retained. If fewer than two pathways were selected, all pathways were used. A multivariate Cox model was then fit using the selected pathways. The composite risk score was defined as:

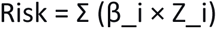

where β_i are the multivariate Cox coefficients. The risk score was tested in TCGA per 1 SD. Coefficients were saved for external validation. A global signature score was also tested as the mean of z-scored pathway predictions in a univariate Cox model.

### 4.9 External validation (GSE65858)

TCGA-derived coefficients were frozen and applied to GSE65858. Raw GSE expression was processed with ssGSEA using the same Hallmark definitions. Features were restricted to the 50 training columns and standardized with the frozen GDSC scaler. Pathway predictions were produced by the same nine models. Predictions were z-scored within each HPV subgroup using GSE-specific mean and SD.

For each HPV subgroup, the risk score was:

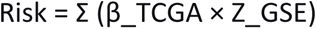

A univariate Cox model on overall survival (OS) was fit, and HR per 1 SD, 95% CI, and p-value were reported. Patients were dichotomized at the median risk score, and Kaplan–Meier curves with log-rank tests were generated.

#### 4.9.1 HPV-stratified replication

HPV+ and HPV− patients in GSE65858 were compared per pathway using a one-sided Mann– Whitney U test (alternative=’greater’). The effect size was computed as 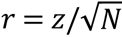, where 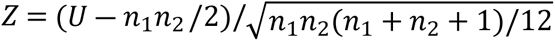. p-values were BH-FDR-corrected across the nine pathways.

#### 4.9.2 HPV-stratified reference-pair replication

Pairwise Spearman correlations were recomputed within each HPV subgroup of GSE65858. A pair was considered replicated only when the GSE ρ matched the TCGA ρ in sign, exceeded |ρ| > 0.2, and had q < 0.05.

### 4.10 Statistical significance thresholds

Unless stated otherwise, significance was defined as BH-FDR-corrected q < 0.05. Network analyses also applied an |ρ| threshold (typically 0.2 or 0.3) to identify biologically meaningful associations. Correlation-coefficient 95% CIs used Fisher’s Z transformation; Hodges–Lehmann difference CIs used a non-parametric bootstrap (1,000 resamples). Coefficient stability was bootstrapped 1,000 times with α and l1_ratio fixed at their full-data values.

### 4.11 AI-assisted writing disclosure

During the preparation of this manuscript, the authors used “DeepSeek Flash 4.1” to summarize the Method and Results sections after they had been written by the authors. They also used the same model throughout the article to improve clarity. All AI-assisted output was reviewed, verified, and edited by the authors. The authors take full responsibility for the content of the publication. No AI tool was used to generate or interpret research data, and no AI tool is listed as an author.

## Data availability

All data used in this study are publicly available. The GDSC fitted dose–response data (GDSC1 and GDSC2) are available from the Genomics of Drug Sensitivity in Cancer project. Compound annotations were obtained from the GDSC screened compounds file (screened_compounds_rel_8.1.csv). The 50 MSigDB Hallmark gene sets were obtained from the Human MSigDB Hs Symbols v2026.1 GMT file. TCGA-HNSC data were obtained from the Genomic Data Commons; the specific files used are listed in the Materials and Methods. GSE65858 expression and survival data were obtained from the Gene Expression Omnibus (GSE65858). GSE103322 single-cell data were obtained from the Gene Expression Omnibus (GSE103322). All other data are within the manuscript and its Supporting Information files.

## Code availability

All author-generated code used to reproduce the analyses and figures is available at https://github.com/yash-0600/HNSCC_pathway_dependency.

## Acknowledgments

None.

## Author contributions

Y.A: Conceptualization, Data curation, Formal analysis, Investigation, Methodology, Software, Validation, Visualization, Writing – original draft, Project administration, Resources, Supervision

A.G: Conceptualization, Methodology, Writing – review & editing

## Competing interests

The authors declare that they have no competing interests.

## Ethics approval and consent to participate

Not applicable

## Consent for publication

All authors have read and approved the final version of the manuscript.

## Funding

The authors received no financial support for the research, authorship, and/or publication of this article.

## Supporting information captions

S1 Table. IC50 missingness analysis.

Mann–Whitney U tests comparing leave-one-drug-out pathway dependency between cell lines with observed vs missing IC50 for each drug with ≥20 tested cell lines. Reported are n per group, medians, p-value, and BH-FDR q-value. Panel outputs also include the p-value distribution and top 20 drugs by proportion of pathways with significant missingness.

S2 Table. Interaction screening: additive vs interaction models.

Per-fold and pathway-level comparison of 5-fold cross-validated R² for the additive (50 Hallmark features) and interaction (50 features + 1,225 pairwise products) ElasticNet models. ΔR² = R²(interaction) − R²(additive). Interactions were retained when ΔR² > 0.01.

S3 Table. Leave-one-pathway-out (LOPO) validation.

For each held-out pathway, all drugs annotated to that pathway were removed from training. Reported are mean Spearman ρ (primary metric) and R² between predicted and held-out dependency scores across 5-fold CV, with the number of held-out drugs and mean train/test cell counts per pathway.

S4 Table. Drug–pathway Spearman correlations.

For each drug with ≥10 measured cell lines, mean LN-IC50 per cell line was correlated with each pathway’s dependency score (leave-one-drug-out for the drug’s own target pathway). Only significant negative correlations (ρ < −0.3, q < 0.05, BH-FDR) are listed, with 95% Fisher Z confidence intervals.

S5 Table. Drug-target validation (High vs Low dependency).

Cell lines were binarised at the 75th percentile of leave-one-drug-out target-pathway dependency, and LN-IC50 values were compared with a one-sided Mann–Whitney U test (alternative = ‘less’). Reported are U, effect size r, Hodges–Lehmann estimate with bootstrap 95% CI, and BH-FDR q-value per drug.

S6 Table. Copy-number validation.

Four a priori hypotheses were tested (EGFR amplification → EGFR; PTEN deletion → PI3K/MTOR; CDKN2A deletion → CELL_CYCLE; CCND1 amplification → CELL_CYCLE). Predicted dependency was compared between Altered and Normal (diploid) tumors using a two-sided Mann–Whitney U test. Reported are n per group, medians, Hodges–Lehmann estimate with bootstrap 95% CI, Z, r, PSet, direction, and BH-FDR q-value.

S7 Table. Mutation-stratified association.

For each gene–pathway pair (TP53 → P53; PIK3CA → PI3K/MTOR; KRAS → MAPK; BRAF → MAPK; EGFR → EGFR; PTEN → PI3K/MTOR), predicted dependency was compared between mutant and wild-type tumors. Variants were classified by functional impact (LoF, Hotspot, Missense, Indel, Other), retaining the most severe class per patient. Only mutation classes with ≥5 mutant samples are reported, with two-sided Mann–Whitney U statistics, Hodges–Lehmann estimate, bootstrap 95% CI, and BH-FDR q-value.

S8 Table. RPPA single-protein validation.

For each protein or phospho-protein mapped to a pathway, Spearman ρ between protein abundance and the matched pathway’s predicted dependency is reported with Fisher Z 95% CI and BH-FDR q-value. A secondary Mann–Whitney U test compares predicted dependency between patients with protein above vs below the median.

S9 Table. HPV multivariate coefficients (TCGA-HNSC).

Per pathway, HPV coefficient from OLS regression of predicted dependency on HPV + age + gender + stage + smoking + treatment score, on complete cases (N = 111; 38 HPV+, 73 HPV−). Continuous covariates were standardized. Reported are β, 95% CI, p-value, and BH-FDR q-value.

S10 Table. CD8⁺ confounding across cohorts.

Spearman ρ between each pathway prediction and the CD8⁺ T-cell infiltration score (mean CD8A, CD8B, GZMA, GZMB, PRF1 expression) in TCGA-HNSC and GSE65858. Cross-cohort concordance of the confounding magnitudes is reported as the Spearman ρ across the nine pathways.

S11 Table. GSE65858 replication of HPV association.

Comparison of predicted dependency between HPV+ (n = 60) and HPV− (n = 196) GSE65858 patients for each pathway using a one-sided Mann–Whitney U test (alternative = ‘greater’). Reported are group sizes, medians, U, Z, rank-biserial effect size r, p-value, and BH-FDR q-value.

S12 Table. CD8⁺-adjusted HPV association (TCGA-HNSC).

Per pathway, HPV coefficient from OLS regression of predicted dependency on HPV + CD8⁺ (N = 114). Reported are β, 95% CI, p-value, and BH-FDR q-value.

S13 Table. Smoking sensitivity for EGFR and MAPK.

Comparison of HPV coefficients between two models fit on the same sample (N = 112): dependency ∼ HPV + CD8⁺ and dependency ∼ HPV + CD8⁺ + smoking. Reported are β and q-values for each model per pathway, and Δβ = β(Model B) − β(Model A).

S14 Table. Phenotype associations with dependency (TCGA-HNSC).

Spearman correlations between each pathway prediction and age, stage, smoking, and alcohol (≥20 paired observations per test). Reported are N, ρ, p-value, and BH-FDR q-value across all tests.

S15 Table. Treatment intensity vs dependency (TCGA-HNSC).

Patients were grouped by treatment intensity: High (≥2 modalities), Low (1), None (0). Mann– Whitney U tests compared High vs. Low and High vs. None per pathway. Reported are group sizes, p-values, and BH-FDR q-values pooled across both comparisons and all pathways.

S16 Table. P53 pathway per-gene mediation.

For each of the 190 Hallmark P53 genes in the TCGA expression matrix, the HPV coefficient was estimated before and after adding the gene to the model (dependency ∼ HPV + gene + age + gender). Reported are Spearman ρ with HPV and with P53 dependency, gene coefficient, attenuation = (β_base − β_adj) / β_base, and BH-FDR q-values for both the gene term and the adjusted HPV term. Ranked by attenuation.

S17 Table. TP53-stratified P53 dependency.

HPV coefficients from OLS models of P53 dependency on HPV + age + gender within All (N = 114), TP53-WT (n = 40), and TP53-mut (n = 4) strata. Reported are stratum size, HPV+ / HPV− counts, β, 95% CI, p-value, and BH-FDR q-value.

S18 Table. p53 transcriptional score by HPV and TP53 status.

Mann–Whitney U comparison of HPV+ vs HPV− tumors for the p53 target score (mean of 13 canonical p53 targets), p53 repressor score (mean of 12 p53-repressed genes), and net p53 activity (targets − repressors), within All, TP53-WT, and TP53-mut strata. Reported are group sizes, medians, U, effect size r, p-value, and BH-FDR q-value.

S19 Table. TCGA co-dependency matrix, all patients.

Pairwise Spearman correlations between the nine predicted pathway dependencies in TCGA-HNSC (N = 565). Reported are ρ, Fisher Z 95% CI, p-value, BH-FDR q-value, and a significance flag (|ρ| > 0.3, q < 0.05).

S20 Table. GSE65858 HPV-stratified co-occurrence.

Pairwise Spearman correlations between the nine predicted pathway dependencies in GSE65858, computed separately within HPV+ (n = 60) and HPV− (n = 196) subgroups. Reported are ρ, Fisher Z 95% CI, p-value, and BH-FDR q-value per subset.

S21 Table. Predicted vs RPPA co-activity.

Cross-layer comparison of pairwise pathway co-activity across the 36 pathway pairs: predicted co-dependency in the RPPA-matched TCGA-HNSC subset (n = 212) vs RPPA directional co-activity (mean z-scored activators − mean z-scored suppressors). Reported are ρ_pred, ρ_rppa, and BH-FDR q-values for each layer.

S22 Table. RPPA candidate dependency nodes.

RPPA proteins classified as direct drug targets, activity readouts, upstream regulators, or pathway suppressors were correlated with matched pathway predicted dependency. Reported are raw Spearman ρ (Analysis A), partial ρ controlling for the other eight pathway dependencies (Analysis B), and BH-FDR q-values. Candidate dependency nodes were defined as direct drug targets with partial ρ > 0.15 and q < 0.10.

S23 Table. Survival analysis (TCGA-HNSC).

Univariate Cox proportional-hazards models of the global dependency signature (mean of z-scored pathway predictions) on progression-free interval, fitted separately within HPV+ and HPV− strata. Reported are N, number of events, hazard ratio per 1 SD, 95% CI, and p-value.

## Notes

### Competing Interest Statement

The authors have declared no competing interest.

